# Unraveling the functional attributes of the language connectome: crucial subnetworks, flexibility and variability

**DOI:** 10.1101/2022.03.31.486594

**Authors:** E. Roger, L. Rodrigues De Almeida, H. Lœvenbruck, M. Perrone-Bertolotti, E. Cousin, JL. Schwartz, P. Perrier, M. Dohen, A. Vilain, P. Baraduc, S. Achard, M. Baciu

**Affiliations:** Univ. Grenoble Alpes, Univ. Savoie Mont Blanc, CNRS, LPNC, 38000 Grenoble, France; Univ. Grenoble Alpes, CNRS, Grenoble-INP, GIPSA-Lab, 38000 Grenoble, France; Univ. Grenoble Alpes, CNRS, Inria, Grenoble INP, LJK, 38000 Grenoble, France

**Keywords:** language, fMRI, connectome, functional connectivity, networks

## Abstract

Language processing is a highly integrative function, intertwining linguistic operations (processing the language code intentionally used for communication) and extra-linguistic processes (e.g., attention monitoring, predictive inference, long-term memory). This synergetic cognitive architecture requires a distributed and specialized neural substrate. Brain systems have mostly been examined at rest. However, task-related functional connectivity provides additional and valuable information about how information is processed when various cognitive states are involved. We gathered thirteen language fMRI tasks in a unique database of one hundred and fifty neurotypical adults (*InLang database*). The tasks were designed to assess a wide range of linguistic processes and subprocesses. From this database, we applied network theory as a computational tool to model the task-related functional connectome of language (LANG). The organization of this data-driven neurocognitive atlas of language is examined at multiple levels, uncovering its major components (or *crucial subnetworks*) and its anatomical and functional correlates. Furthermore, we estimate its reconfiguration as a function of linguistic demand (*flexibility*), or several factors such as age or gender (*variability*). By accounting for the multifaceted nature of language and modulating factors, this study can contribute to enrich and refine existing neurocognitive models of language. The LANG atlas can also be considered as a reference for comparative or clinical studies, involving a variety of patients and conditions.

## 1 Introduction

Language is optimized for human communication. It is an efficient vector of information transmission, shaped under cultural and environmental constraints to meet physical, technological and social needs (e.g., Kirby et al., 2015; Lupyan & Dale, 2016; Millikan, 2005; Scott-Phillips, 2015; Tamariz & Kirby, 2016). Language is also adapted to thinking and interpretation, playing a scaffolding role in cognition (e.g., Carruthers, 2002; Chomsky, 2014; Clark, 2006; Jackendoff, 1996; Reboul, 2015). Finally, language plays a crucial role in meta-cognition including self-evaluation, self-regulation and autonoetic consciousness (Alderson-Day & Fernyhough, 2015; Perrone-Bertolotti et al., 2014).

In order to combine both effectiveness (Gibson et al. 2019) and utility (Jaeger & Tily, 2011) of language production and comprehension, several essential abilities are required. The first one, is the combinatory skill (Boer et al., 2012; Friederici et al., 2017; Zuidema & de Boer, 2018). Language is compositional and recursive, implying specialized processing of intra-linguistic aspects (i.e., an aptitude to handle various combinatorics, perceptive, syntactic or semantic/conceptual; Pylkkänen, 2019). A second important ability relates to multisensory integration which arguably facilitates spoken communication and enhances speech intelligibility (Chandrasekaran et al., 2009; Ghazanfar & Schroeder, 2006; Luo et al., 2010; Noppeney et al., 2008; Schroeder & Foxe, 2005; Schwartz et al., 2004; Sumby & Pollack, 1954). Beyond the multisensory facilitation (or low-level multimodal integration; Holler & Levinson, 2019), high level cognitive abilities related to top-down multimodal mechanisms have also been emphasized. A shared understanding, a relevant and contextually-adapted discourse, requires aligning the partners’ representations, considering shared knowledge, past experiences, or even making assumptions about the other’s perspectives. Establishing “common ground” between conversational partners (Clark & Marshall, 1981) relies on a wide range of “high-level” cognitive functions such as working memory (resonance-based theory of common ground, Horton, 2007), long term memory (Brown-Schmidt & Duff, 2016), theory of mind or mentalizing (Vanlangendonck et al., 2018) processes. Language use is therefore adapted online, enabling communication in a range of environmental and social contexts, meeting various cognitive demands and individual metacognitive needs. Given the pressure exerted by communication, cognitive and metacognitive demands, language has evolved as an adaptive and complex system, which requires taking into account external (context) and internal (individual needs and goals) signals, while processing both intra- and extra-linguistic signals (e.g. Holler & Levinson 2019, for a multimodal language-in-situ framework).

How are these various abilities integrated in order to sustain language functions and how are they implemented in the brain? The exploration of brain networks and the unique lens it provides for understanding cognitive function has become an important part of the cognitive neuroscience landscape (Fornito et al., 2013). Brain network descriptions have revealed that the brain is organized as a “small-world” network (Achard & Bullmore, 2007), favoring optimization of information transfer (Laughlin & Sejnowski, 2003). This organization is characterized by a balance between segregation and integration, that is by short communication paths creating specialized subsystems (segregation), whose interconnectivity is coordinated by distant highly connected brain regions (integration; Heuvel & Sporns, 2013). Functionally, local systems or highly connected “modules” for the processing of information in a given modality (visual, auditory, etc.) are linked together by sparse and specific fiber paths over long distances, according to a connectivity principle of “local richness and long-range sparseness” (Pulvermüller, 2018). This organization allows efficient serial, parallel and distributed brain activity (Herbet & Duffau, 2020). Integrative areas (sometimes referred to as connector hubs), at the interface between local systems, are particularly important for the multimodal neural integration of information (Cocchi et al., 2013; Fornito et al., 2015; van den Heuvel & Sporns, 2013). With regard to the language circuitry, integration/segregation subsystems and specific connector hubs have been previously identified for both language production and comprehension (e.g., Friederici, 2012; Hagoort, 2016; Hertrich et al., 2020; Roger et al., 2022). However, language is multi-faceted and a comprehensive analysis of the functional properties of the language connectome in a broader framework would contribute to a more accurate description.

In an attempt to fill this gap, in this study, we propose to examine the functional attributes of language at the brain level through an integrative perspective, mixing several linguistic tasks explored in the light of graph theory. This research is naturally framed within the substantial legacy of the study of language and its brain foundations (since the beginnings of modern neurology) whose growing and diverse observations have been accompanied by the evolution of investigative techniques. In the past decades, many authors have highlighted brain function and structure associated with language through theoretical neurocognitive models (e.g., Duffau et al., 2014; Friederici et al., 2017; Hagoort, 2016, 2019; Hickok & Poeppel, 2007; Indefrey, 2011; Levelt, 1989; Price, 2012; Rauschecker & Scott, 2009). However, this study, which is in direct continuation with past legacy, adds further value by investigating functional cerebral connectivity (FC) based on task data. The anatomo-functional substrates associated with language are indeed highly task-dependent (Hickok & Poeppel, 2000).

The task-based FC analyses presented here rely on an fMRI database compiling a broad spectrum of language-related tasks (*InLang* database: doi.org/10.5281/zenodo.6402396). More precisely, *InLang* is composed of thirteen language tasks, performed cross-sectionally by 150 right-handed neurotypical adults. The database is unique in that it covers a broad spectrum of language features: semantic and conceptual processing, decoding (phonology, sound), lexico-syntactic formulation (production), dialogality (social aspects of language), monitoring of self and others, and unintentional speech (Figure 1A; Appendix S1). Such a database is essential to uncover the functional architecture of the multifaceted language processes in an integrative approach. It makes it possible to modelled a comprehensive connectomic atlas of language and to explore its fundamental properties in depth. To this end, we analyzed task connectomes using graph metrics applied at multiple scales, which allowed us to expose: (1) the overarching FC profile of different language tasks and latent subprocesses; (2) the architecture of the general language connectome (LANG); (3) the functional roles of crucial language subnetworks and brain regions; (4) the anatomo-functional correlates; and (5) the flexibility and variability exerted on the LANG connectome. Figure 1 provides an overview of the *InLang* database and the methodology used to address these 5 main axes.

**Figure 1:**
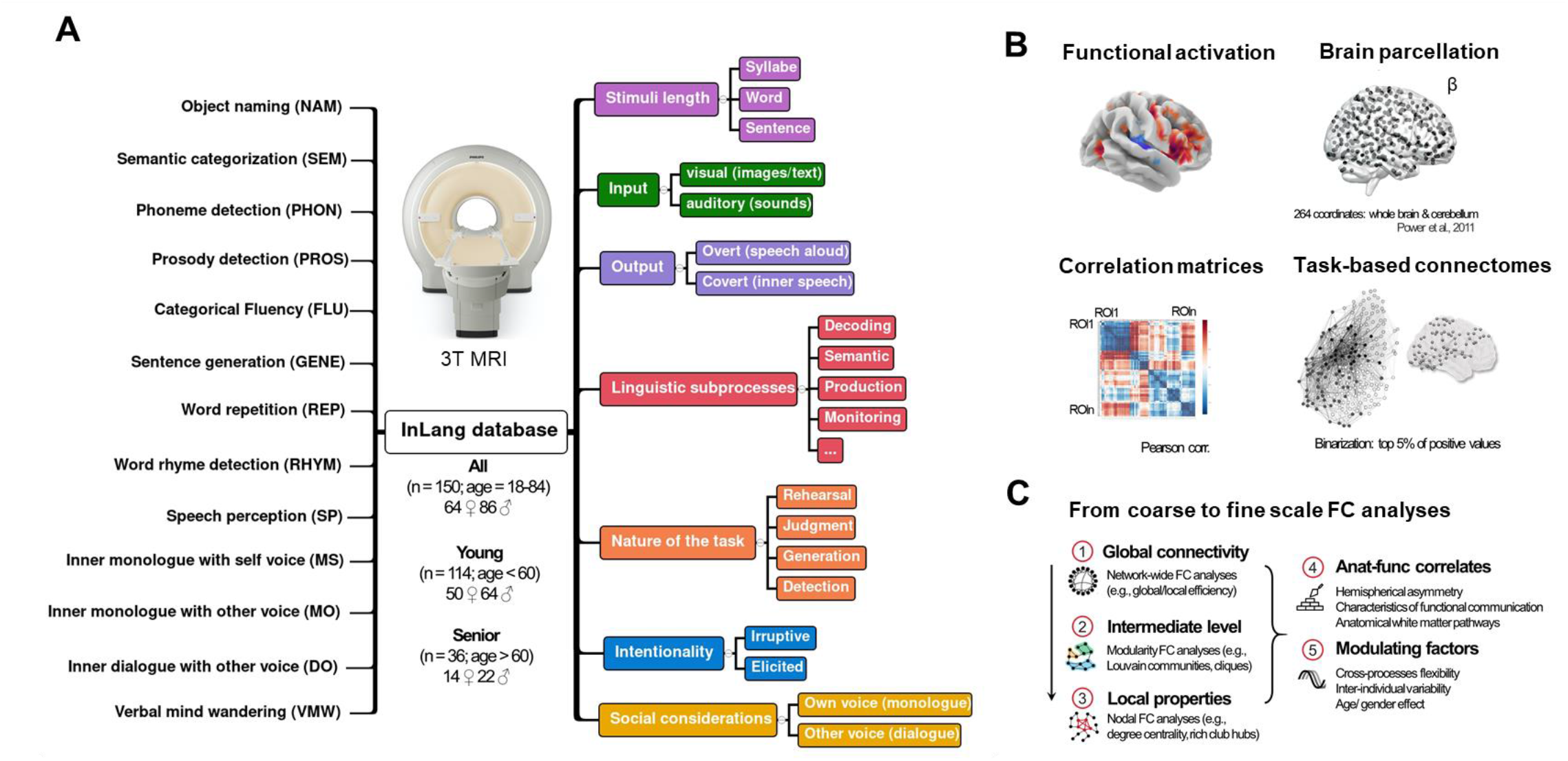
Overview of the *InLang* database and methodological outlines. **A.** The *InLang* database: the thirteen language tasks and the main dimensions manipulated by the protocols. The protocols have been previously published and the MRI data have been acquired between 2010 and 2019. *InLang* gathers, in a unique database, a cross-sectional cohort of 150 different healthy individuals and 359 functional scans (see Materials and Method Section 5.1 and Appendix S1 for more details about tasks and protocols). Table S2 contains the subjects’ characteristics, by tasks. **B.** Brief summary of the steps performed to obtain the task connectomes. For a given task, we extracted the beta values from the individual functional activation maps, on a parcellation covering the whole brain (Power et al., 2011). The beta values were then used to compute the task-specific connectivity matrix (correlation matrix). The same procedure was repeated for all tasks to obtain the respective functional matrices and connectomes. **C.** Outline of the multi-level statistical analyses performed on functional connectivity (FC) measures (i.e., graph theory parameters) to address 5 main axes.

## 2 Results

### 2.1. Towards a task-based connectomic atlas of language

#### 2.1.1. Global profile of the tasks and latent subprocesses

Data-driven clustering, based on the similarity between the global FC profile of the 13 tasks (i.e., a profile combining the global efficiency: *E_glob_*, the local efficiency: *E_loc_*, the mean geodesic distance: *d̄*, and the total number of network nodes: N; Figure 2A), reveals an optimal 5-cluster solution. This solution is consistent with that obtained based on BOLD functional activations (Appendix S1; Figure S1). The internal composition of the five task groups was used to label them according to the underlying language subprocess that might be primarily involved (Figure 2A), namely: G1 = MONITORING (MS, MO, DO tasks); G2 = DECODING (PHON, RHYM and PROS tasks); G3 = SEMANTIC (SEM and SP tasks); G4 = PRODUCTION (NAM, FLU, GENE, and REP tasks); G5 = WANDERING (VMW task). Indeed, the monologal and dialogal inner speech with own and other voice (MS, MO, DO) mainly engage MONITORING processes (inner voice *control*). Phoneme detection (PHON), rhyme judgment (RHYM) and prosodic detection (PROS) first involve phonology and/or prosody DECODING (sound *control*). Semantic categorization (SEM) and speech perception (SP) respectively engage word and sentence comprehension. They both primarily require SEMANTIC processing (conceptual *knowledge*). Object naming (NAM), categorical fluency (FLU), sentence generation (GENE), word repetition (REP) rely on lexical/lexico-syntactic formulation or word PRODUCTION (conceptual *knowledge*). Finally, verbal mind wandering (VMW) involves spontaneous speech production underpinned by introspective WANDERING processes (or *unintentional* thought). Details of the tasks and the language subprocesses theoretically and primarily targeted according to statistical fMRI contrasts performed are provided in Appendix S1 (see also the summary Table S1).

**Figure 2:**
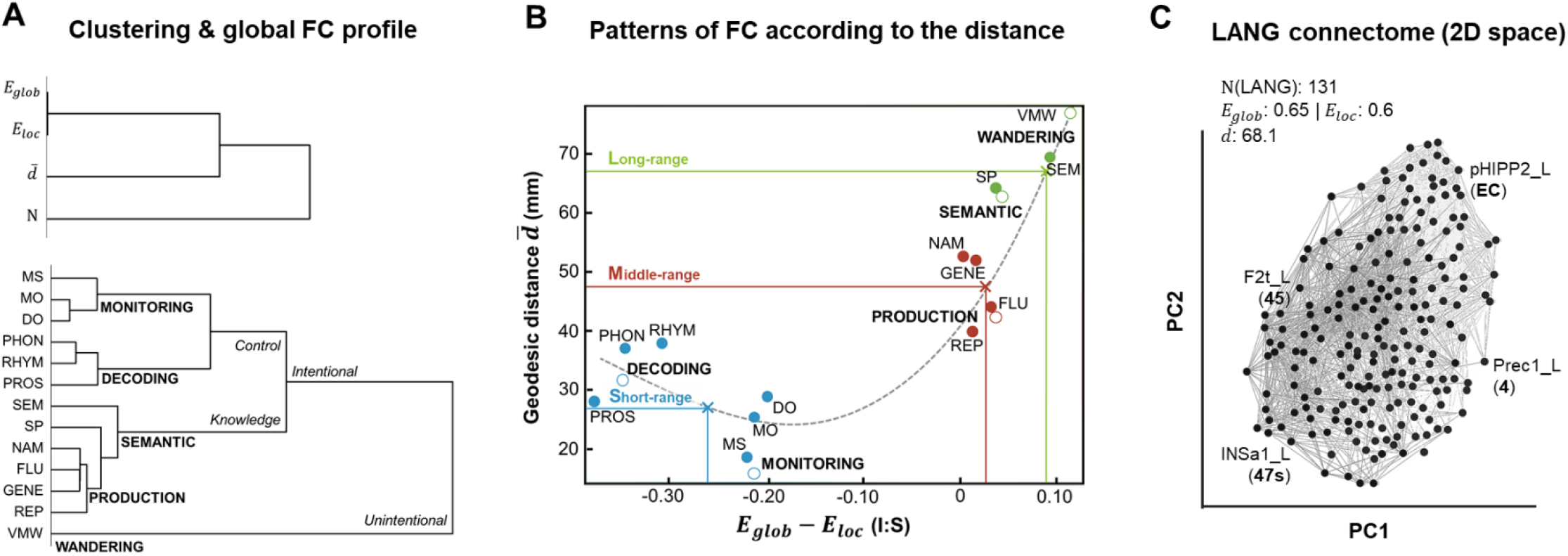
Global connectomic profiles of the tasks, the subprocesses and the LANG connectome. **A.** Hierarchical clustering of parameters used to define global FC profiles (*E_glob_* = global efficiency, *E_loc_* = local efficiency, *d̄* = mean geodesic distance; N = total number of nodes; top); and of individual tasks, to cluster them into groups of underlying subprocesses (bottom). See also Appendix S1 for the rationale of the subprocesses labels (Table S1) and for the clustering applied to BOLD functional activations (Figure S1). Table S3 in Appendix S2 summarizes the global measures for each task and subprocess. **B.** Non-linear significant relationship between the integration/segregation balance (I:S) and the average geodesic distance of the network’s functional connections, for the different tasks and subprocesses. 3 types of connectivity are distinguished according to the I:S/geodesic distance profile (short-range, middle-range, and long-range). The colored lines come from the centroids estimated from the observed data (at the level of brain region) in relation to the regression polynomial curve. **C.** Global topology of the LANG task-based connectome. The 131 regions of interest (ROIs) of LANG (Power et al., 2011; LH = 61%, RH = 35%, CER = 4%) are projected here in a reduced two-dimensional space (PCA layout) and anatomo-functional labels of randomly selected LANG ROIs are shown for illustrative purposes only (Table S4 includes their full name).

Interestingly, language tasks and subprocesses can also be grouped according to the physical distance of their functional connectivity within the brain. By contrasting the integration/segregation balance (I:S) with the mean geodesic distance *d̄*, we observe a significant relationship between the two parameters (*r* = 0.8, *p* < .001; Figure 2B), bringing out a gradual organization of tasks and subprocesses according to 3 canonical profiles of average connectivity: C1 = long-range connections; C2 = middle-range connections; C3 = short-range connections (Figure 2B). The more segregated rather than integrated the networks are (i.e.,negative difference, in favor of *E_loc_*), the shorter the physical internodal distance (short-distance functional connections, as for the control tasks of language involving DECODING and MONITORING subprocesses). Conversely, the more integrated rather than segregated the networks are (positive difference, in favor of *E_glob_*), the longer the physical internodal distance (long-range functional connectivity, as for the WANDERING and SEMANTIC task groups).

#### 2.1.2. Global topology of the general LANG connectome

After excluding irrelevant functional connections (see Material & Methods, Section 5.2.), LANG is composed of 131 non-isolated regions of interest (ROIs; Power’s parcellation: Power et al. 2011), distributed over the two hemispheres (nLH = 80; nRH = 46) and the cerebellum (nCER=5). Connectivity between LANG ROIs appears balanced between integration and segregation (I:S = 0.049), associated with a rather long-range connectivity profile (*d̄*= 68.1). Table S3 (Appendix S2) summarizes the global network properties of the tasks, of the subprocesses and of the general LANG connectome. Figure 2C shows the LANG connectome as a graph projected into a reduced two-dimensional space.

#### 2.1.3. LANG partition and hubs (intermediate and local scale)

Community-based detection applied to the LANG connectome defines 4 distinct components (or functional subnetworks, called Nets; see Figure 3A, for projection into a reduced space). Figure 3B shows the mapping of the Nets onto the brain and cerebellum templates. Figure 3C highlights their internal composition in terms of discrete intrinsic networks (as previously characterized by Ji et al. 2019; Cole-Anticevic Brain-wide Network Partition: CAB-NP).

**Figure 3:**
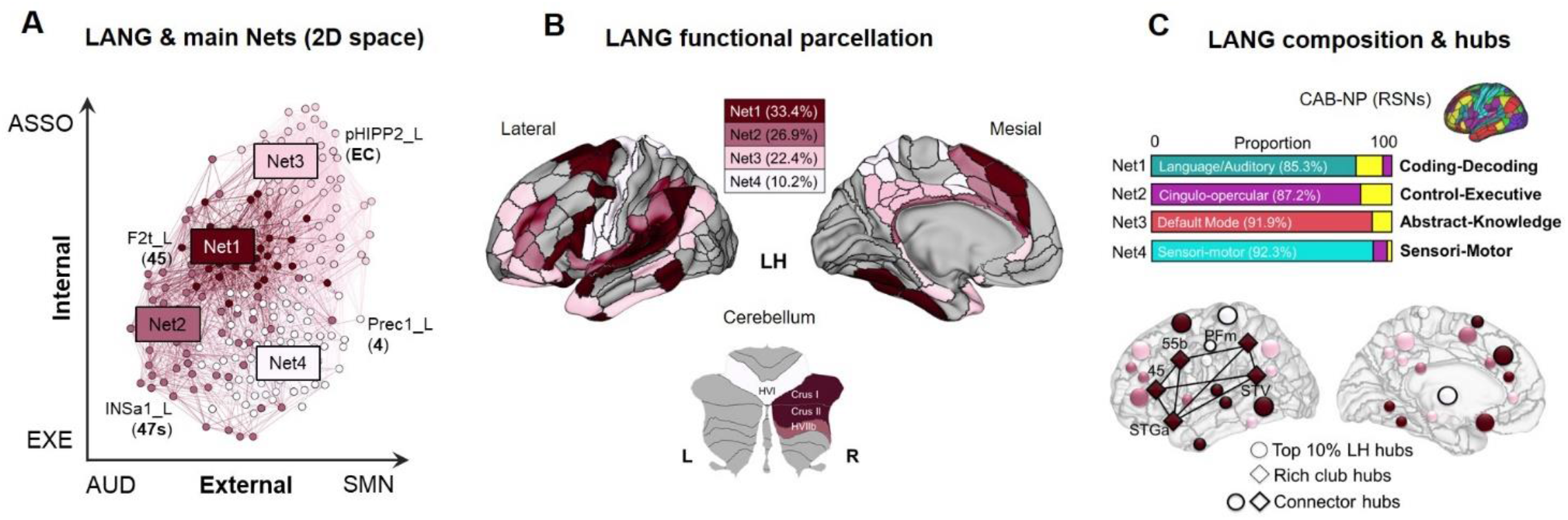
Global, intermediate and local connectomic features of the LANG connectome. **A.** Four main components identified within the LANG network (optimal partition, Louvain method). The components (Nets) are displayed on the connectome projected in the reduced 2D space. The number of ROIs per Nets (Power ROIs; including both hemispheres and cerebellum) is distributed as follows: Net1 = 54 (41.2%), Net2 = 28 (21.4%), Net3= 26 (19.8%), Net4 = 23 (17.6%). ASSO = associative; EXE = executive; AUD = auditory; SMN = sensorimotor. **B.** Illustration of the LANG connectome and its 4-Net functional subdivision on the parcelled brain. Distribution on a multimodal parcellation of the brain (HCP_MMP1.0; Glasser et al. 2016) and cerebellum (SUIT; Diedrichsen et al. 2009). Only the left hemisphere (LH) is represented here (see Appendix S2 for a complete representation of the connectomic atlas) **C.** Intrinsic functional composition of LANG Nets (top), in accordance with the resting state networks (RSNs) proposed by Ji and collaborators (Cole-Anticevic brain-wide network partition; CAB-NP; Ji et al., 2019) on the same template (HCP_MMP1.0 borders). Nodal properties of LANG (bottom), showing the distribution of the main hubs, connector hubs as well as the regions belonging to the maximal clique (complete subgraph; diamond). Only the LH is represented here.

Considering the composition, Net1 could correspond to the core component of language, engaged in the coding-decoding of linguistic signals of multiple nature: e.g., acoustic, syntactic, conceptual, articulatory (*Coding-Decoding system*). Net2 is represented by executive-attentional functional networks (*Control-Executive system*). Net3 is mainly composed of regions of the default mode network (DMN) known to be involved in high-level cognitive abstraction and can thus be regarded as a “conceptual” knowledge network (*Abstract-Knowledge system*). Finally, Net4 involves a large majority of perceptual and motor brain areas, suggesting that it is the “*Sensori-Motor*” *system* of language. A supported argument and in-depth discussion of the putative functional roles of these LANG Nets is raised in the discussion section (Section 5).

The Nets’ composition, coupled with their topological organization in the reduced space, provides evidence for the possible meaning of the 2 main axes (i.e., the principal components; Figure 3A). PC1 extends from auditory to sensorimotor components of language and may reflect the axis of externally oriented cognition (from verbal-specific to domain-general somatosensory systems). PC2 progressively involves control executive regions to semantic associative regions and may represent the axis of high-level internal cognition associated with language.

Interestingly, the core Net1 is located at the crossroads of these two internal-external axes. Moreover, Net1 is the component with the highest portion of connector nodes (Net1= 40.7%, distributed in both hemispheres; Table S4, Appendix S2) reflecting a high capacity to integrate information from regions belonging to the same network (intra-FC), as well as to other specialized networks (inter-FC; high *zi*, high *Pi* class). Net1 also exhibits a “rich club” organization (from a rich club regime of k >17 to k < 26; Φnorm(k) >1, p < .001, 10.000 permutations). Restricting to the level of k where the strongest rich club effect was observed (k=24), we found a set of 5 left perisylvian hubs constituting the “rich-club” of Net1 (areas: STGa, 45, 55b, PFm, STV; Figure 3C). These nodes also form the maximal Net1 clique (i.e., the maximal complete subgraph; ω(LANG/Net1) = 5).

Table S2 includes information about the LANG modules and hubs for each region. Appendix S2 presents the LANG connectomic atlas including the right hemisphere (RH), as well as details of its components.

### 2.2. Properties of the LANG connectomic atlas

#### 2.2.1. Functional correlates

On average, the LANG’s FC laterality index indicates a slight LH predominance (LI(LANG)= +0.23), but hemispheric asymmetry is variable across the Nets. The proportion of nodes that are more strongly connected is higher in LH for Net1 (LI(Net1) = +0.41) than for the other Nets (Figure 4A). By comparison, the FC of the nodes belonging to the “sensory-motor component” are bilaterally distributed (LI(Net4) = −0.12). Overall, the FC asymmetry of LANG Nets (from bilateral to LH) is arranged along the following gradient: Net4 < Net3 < Net2 < Net1.

**Figure 4:**
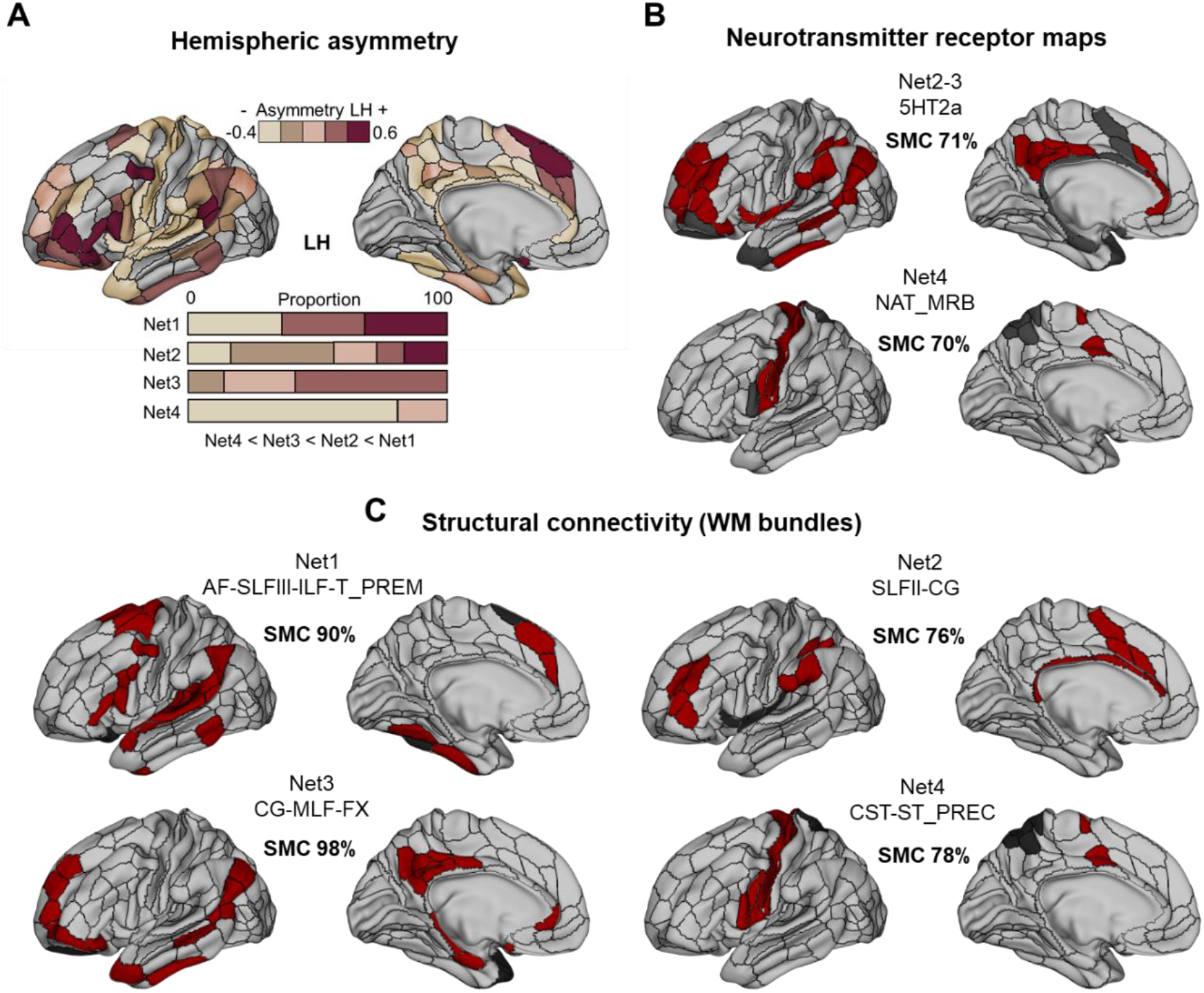
Functional attributes and structural underpinnings of the LANG connectome. **A.** Asymmetry of FC estimated on LANG ROIs and their distribution as a function of Nets. Left hemispheric FC dominance can be considered if IL > +0.2 (see Method section). **B.** Spatial concordance with neurotransmitter receptors maps (receptomes) of: serotonergic (5HT2a [F18]altanserin PET; Savli et al. 2012) and catecholaminergic/noradrenergic (NET (S,S)-[(11)C]O-methylreboxetine (MRB) PET; Hesse et al. 2017) pathways. All the neurotransmitters’ maps implemented in JuSpace (Dukart et al., 2021) were tested, but only spatial matches considered sufficient are shown here (SMC > 0.67; see Method section). LANG regions with significant coverage (>40% of overlap) are in red: those with no or insufficient coverage (<40%) are in gray. **C.** Structural concordance with large white matter (WM) bundle terminations provided by TractSeg (Wasserthal et al., 2018). Only the best bundles combinations allowing for the highest match are displayed here. The red/gray color code corresponds to the same definition as for Panel B.

In addition, some LANG Nets are spatially congruent with the mapping of neurotransmitter receptor pathways. In particular, the LH nodes of Net2 and Net3 show a high spatial matching with the serotonin receptors 5HT2a (SMC Net2/5HT2a = 0.68; SMC Net3/5HT2a = 0.74). Those of Net4 overlapped with the noradrenergic transporters NAT_MRB (SMC Net4/NAT_MRB = 0.7). Figure 4B shows the distribution of the LANG ROIs that match (or do not match) with the PET receptors (see also Figure S3 in Appendix S2).

#### 2.2.2. Structural correlates

The endings of some large white matter (WM) bundles are spatially concordant with the LANG Nets (Figure 4C; Figure S2 in Appendix S2). The best overlap between the bundles and Net1 (LH ROIs) is obtained by combining the ending masks of the left arcuate fascicle (AF), superior longitudinal fascicle branch III (SLFIII), inferior longitudinal fascicle (ILF) and the thalamo-premotor (T_PREM) projections (SMC Net1/AF-SLFIII-ILF-T_PREM complex = 0.9). The concordance rate increases to 92% when the ending masks of the middle cerebellar peduncle (MCP) and the cerebellar ROIs of Net1 are included. At a more restricted level, the unique contribution of the left AF provides a high spatial concordance with the Net1 lateral LH nodes (SMC Net1/AF = 0.72). Regarding Net2 (LH ROIs), the best matching is reached with the combination of the left superior longitudinal fascicle branch II (SLF-II) and the cingulum (CG) bundle (SMC Net2/SLFII-CG = 0.76). The concordance between the SLFII individually taken and the lateral LH Net2 ROIs is close to 70% agreement (SMC Net2/SLFII = 0.68). Net3 (LH ROIs) has almost complete coverage when considering the combination of CG, the middle longitudinal fascicle (MLF) and the fornix (FX; SMC Net3/CG-MLF-FX = 0.98). Finally, Net4 (LH ROIs) is spatially well covered by the combination of the cortico-spinal (CST) and the striato-precentral (ST_PREC) tracts (SMC Net4/CST-ST_PREC = 0.78).

#### 2.2.3. Flexibility and variability

The module assignments of LANG ROIs vary according to the linguistic subprocesses involved. We have calculated the flexibility coefficients to capture the FC versatility of the ROIs engaged in the different Nets, depending on the subprocess at work. The average flexibility coefficient (F) is rather low for Net1 (F Net1 = 0.21) and Net4 (F Net2 = 0.32); while those for Net2 and Net3 are twice as high (F Net2 = 0.55; F Net3 = 0.64). Ordering LANG networks according to their functional versatility yields: Net1 < Net4 < Net2 < Net3; from invariant to highly flexible (Figure 5A).

**Figure 5:**
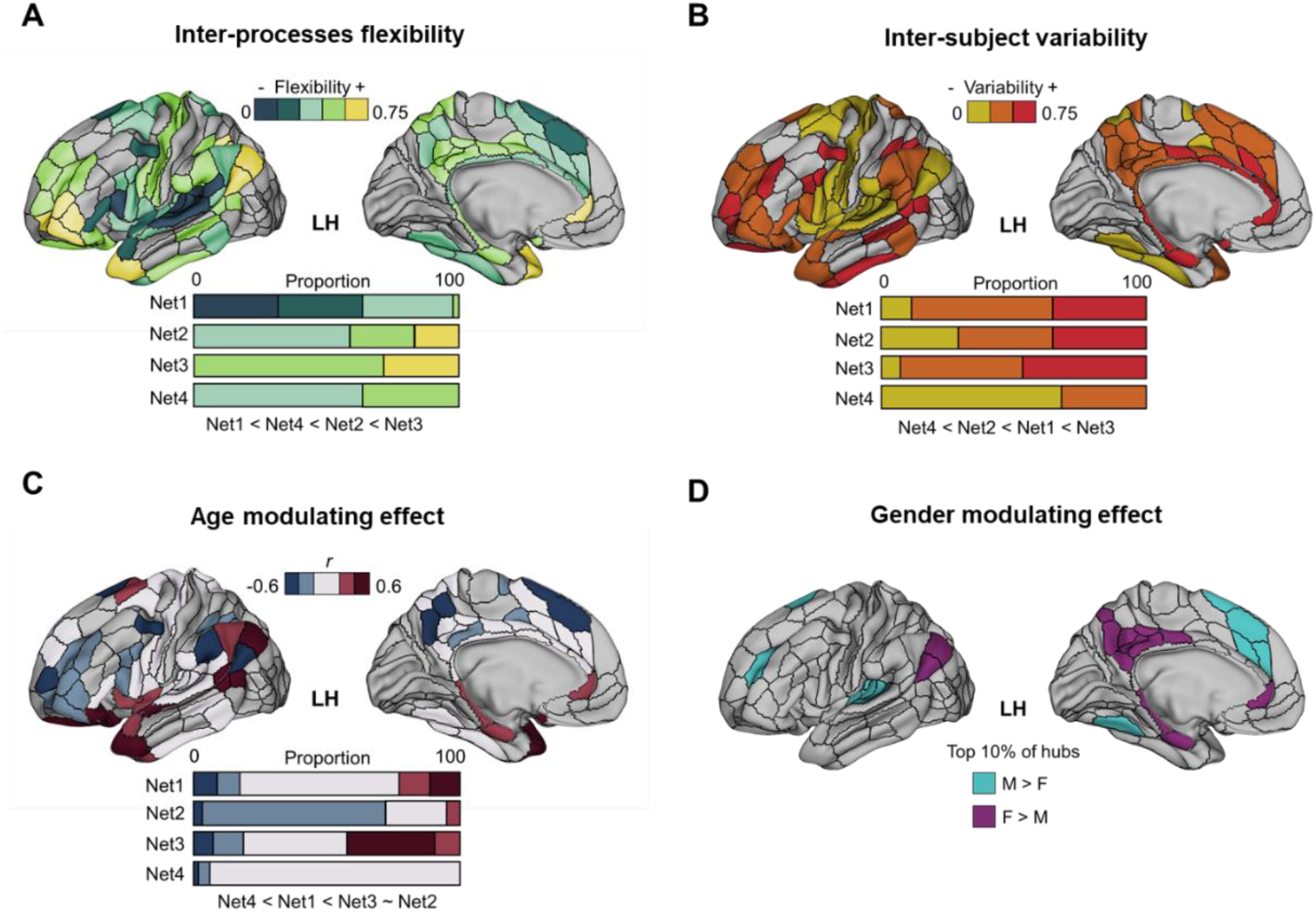
Variability of the functional attributes of the LANG connectome. **A.** Variability induced by the linguistic demand (i.e., the linguistic sub-process involved in the task). Representation of the flexibility score of each of the LANG ROIs as well as their distribution in each of the Nets. **B.** Inter-individual variability. Representation of the betas z-scores estimated on the LANG ROIs for all subjects (young) as well as their distribution in each of the Nets. **C.** Variability determined by age. Illustration of the correlation coefficients between age and ROI DCs, as well as their distribution in each of the Nets. The correlations were performed on the naming task (NAM) that includes healthy participants over a wide age range: 82 subjects aged 18 to 84 years. **D.** Variability induced by gender. Top 10% of the most different LANG ROIs between males (M) and females (F; self-reported gender). In cyan, the top 10% of ROIs where nodal connectivity (DC) is higher in males compared to females. In magenta, the top 10% of regions where nodal connectivity is comparatively higher in females than in males.

Although modest, there is also some inter-individual variability in FC when individuals perform the language tasks (Figure 5B). We find the highest inter-subject variability on Net3, but the variance remains low on average (mean z score = 0.56). FC variability between participants is more visible at the regional level than at the network scale. In addition, we found a high matching coefficient between the “universal language network” (ULN; as proposed by Ayyash et al., (2021) and the lateral LH ROIs of Net1 (SMC Net1/ULN = 0.84; Appendix S2), suggesting some between-individual and cross-cultural consistency in key language network involvement.

However, the LANG connectome undergoes changes with age (Figure 5D). We observe both positive and negative correlations between age and degree centralities (DCs). Net3 and Net2 are the components showing the most important modulations with age. More specifically, ROIs of Net2 are negatively correlated with age (mean r = −0.39); while ROIs of Net3 are, on average, positively correlated with age (mean r = 0.31). Thus, the older the individuals, the less functionally connected the Net2 regions are (decrease in functional hubs for this network in LANG). By contrast, the Net3 regions tend to be more strongly interconnected in LANG with age.

Finally, gender also modulates LANG connectivity. The strongest LANG ROIs for males compared to females (M > F) in terms of DCs are distributed between Net1 (61.54%) and Net2 (38.46%) in LH. The strongest LANG ROIs for females compared to males (F > M) are practically all located in Net3 (92.3%). Figure 5C shows the LANG ROIs with the most divergent FC.

## 3 Discussion

The main objective of this study was to provide an in-depth, multi-scale view of the organization of brain function associated with language from a connectomic perspective. We leveraged an extensive fMRI database of multi-paradigm language tasks (*InLang* database) and we applied a state-of-the-art functional connectivity (FC) methodology that provides unique insights on brain networks. The central finding of this research is that the general language connectome can be objectively partitioned into four main non-overlapping subnetworks (referred to as “Nets”), possessing distinctive and marked features. Table 1 provides a complete overview of these Nets.

**Table 1:** Summary of the properties associated with each of the language networks.

| LANG Nets | Composition & main features | Functional attributes | Anatomical underpinnings |
| --- | --- | --- | --- |
| **Net1**<br>*Coding-Decoding System* | - Language/auditory-verbal component (RSNs: LANG/AUD)<br>- Strongly interconnected (perisylvian rich-club)<br>- At the crossroads of the other Nets (Core) | - Systematically engaged in language tasks (unflexible)<br>- Compatible with the “universal language network”<br>- LH lateralized | - Supported by a complex of long-range fascicles (AF-SLFIII-ILF-ST_PREM; lateral part: AF +++) |
| **Net2**<br>*Control-Executive System*<br>**Net3**<br>*Abstract-Knowledge System* | - Associative regions involved in high-level cognition (RSNs Net2: CON/FPN; Net3: DMN) | - Flexible according to the linguistic subprocesses involved (versatile)<br>- Age-sensitive (Net2: DC ↓ with age; Net3: DC ↑ with age)<br>- Serotonergic pathway | - Long-range associative WM fascicles (Net2: SLFII-CG; Net3: MLF-CG-FX) |
| **Net4**<br>*Sensori-Motor System* | - “Grounded” cognition (RSN: SMN) | - Globally engaged in language tasks<br>- FC bilaterally distributed<br>- Resistant to aging<br>- Noradrenergic pathway | - Coupled with projection fibers (CST-ST_PREC) |

The most extensive subnetwork, Net1, corresponds to a “specialized” language system shaped for the encoding-decoding of auditory-verbal signals. Indeed, Net1 consists primarily of areas belonging to the intrinsic networks previously designated as “auditory” and “language” (CAB-NP RSNs: Ji et al. 2019, Figure 3C). It indeed includes a set of both primary, secondary and associative areas, previously noted as specialized for language (e.g., Labache et al. 2019; Price 2012). In particular, a subset of Net1 composed of key brain regions densely interconnected to each other, forms a critical set for information integration and communication during language tasks (Figure 3C). These core network structures are inscribed in the left perisylvian zone, namely: the anterior part of the superior temporal gyrus (STGa), the posterior part of the pars triangularis of the inferior frontal gyrus (pIFG, 45/44), posterior middle frontal gyrus/premotor cortex (pMFG, 55b area), inferior parietal cortex, supramarginal gyrus (PF/PFm,), temporo-parieto-occipital junction (STV/TPOJ1). Our analyses to determine the functional role of brain regions within networks have identified them all as “connector” hubs, which is consistent with previous observations (e.g., IFG/TPJ/STG: Goucha, Zaccarella, et Friederici 2017; Hagoort 2016; PF/SMG: Braga et al. 2013; pMFG, 55b: Hazem et al. 2021). Connector hubs are most likely to be located at the contact points of several white matter (WM) fascicles, actively supporting long-distance information transport and processing (e.g., at the AF/SLF convergence areas for the IFG and TPJ; Roger et al. 2022). We have indeed observed a strong matching with the AF endpoints (whose involvement in language has been widely and more directly reported; e.g., Forkel et al. 2022 for a meta-analysis) and the lateral perisylvian part of Net1. But more than a one-to-one relationship between structure and function, a combination of various WM bundle terminations seems to underlie the entire network (Figure 4C). In addition, and beyond the clique, Net1 embeds other connector areas, some of which in the right hemisphere, others in the basal ganglia (the anterior parts of the thalamus and the putamen) or in the right cerebellum (Crus I) as well (Table S4, Appendix S2). Large cortico-basal ganglia-cerebellar loops would be involved during language tasks, supporting a substantial role of the subcortical structures in high-level cognition including language (Murphy et al., 2022).

Individually, brain areas have their own anatomical and microstructural properties (cytoarchitectonic features, Zilles et Amunts 2010) and may thus be biased – under normal conditions – to respond efficiently and preferentially to certain types of input. They can be tuned for functional selectivity to linguistic phonological, syntactic, lexical or even semantic units (Friederici 2011). However, the underlying computational processing (i.e., the functional role) of regions belonging to the same Net could be deeply similar. Computational building blocks (called primitives: Poeppel 2012; elementary linguistic operations: Hagoort 2019; or neural operations: Buzsáki 2020) of Net1 could imply here the segmentation-fusion of the linguistic signal, yielding the generation of a verbal information stream of increasing and ordered complexity (Zaccarella & Friederici, 2015). Multiple combinatorial operations of language on different linguistic representations have already been reported (e.g., the combinatorial network of language of Pylkkänen 2019). Net1 and its constituents could represent the foundation of these combinatorics in task. The modularity analysis we applied on multiple language tasks would indeed have captured a common “language combinatorial” computational mechanism for Net1, making this network a cornerstone of a “language-specialized” encoding-decoding system (Hagoort, 2017).

Consistent with a central system, Net1, is topologically situated at the interface of the other components, between an internally and externally oriented cognition (Figure 3A). Moreover, Net1 was found to be a globally inflexible (unchanged) configuration regardless of task and linguistic demand (Figure 4A). It also appears spatially consistent with the “universal language network” proposed by Ayyash et al. 2021, as an invariant, cross-cultural, functional language network (see Appendix S2). A number of universals of language (apart from the “universal grammar”; Chomsky 1995, which is debated) have been reported (Coupé et al., 2019) and concern both semantics (Gibson et al., 2017), syntax (Futrell et al., 2015) or even pragmatics (Piantadosi et al., 2011). The constraints applied to shape languages seem to follow common rules of optimization of coding and information transfer towards a fundamental principle of efficiency. The functional selectivity of Net1 regions is likely to be inherited from our ancestors and to be part of a language-ready brain (Boeckx & Benítez-Burraco, 2014). They are also supported by a specific brain architecture already present in children (Friederici, 2017) whose functional connectivity is genetically encoded (Mekki et al. 2022, for the genetic regulation specifically involved in the perceptual-motor and semantic pathways of language).

At the boundaries of Net1, we also detected two networks that are integral parts of the general LANG connectome (Nets2-3). Net2 is dominated by intrinsic attentional and executive control networks (cingulo-opercular and fronto-parietal networks; Figure 3C). First, the cingulo-opercular network (CON) is a superordinate system encompassing the salience network (Ji et al., 2019), involved in external-signal-driven attentional control or top-down, “exogenous” redirection of attention (Matthen, 2005). The specialization of such a network in the active controlled integration of exteroceptive information may lead to the provision of relevant information in working memory (Parr & Friston, 2017) in order to construct an internal representation of the external world that is relevant to the individual at a specific time. Second, the fronto-parietal network [FPN; close to the Multiple Demand Network (MDN): Smith et al., 2021, or to the Central Executive Network (CEN); Doucet et al., 2019] is a network involved in all processing requiring controlled attention directed toward internal cues and goals. This network operates for endogenous and top-down attentional redirection (Perrone-Bertolotti et al., 2020) and is engaged in verbal working memory and “fluid” cognition (Assem et al., 2020). Overall, Net2 is a controlled, executive language system that captures both endogenous and exogenous attentional aspects. Net3, on the other hand, is almost exclusively composed of DMN regions (Figure 3C). At rest, the default state is thought to be involved in “random episodic silent thought” promoting creativity (Andreasen, 2011). Task-based studies have shown its involvement in natural language processing (Simony et al., 2016). As a foundation of the episodic-semantic memory spectrum (and more broadly language-memory; Roger et al. 2022), the DMN is a multimodal experiential system (Xu et al., 2017) that fosters resonance and binding between environmental features and those derived from similar prior knowledge and states (Binder & Desai, 2011; Constantinescu et al., 2016). For these reasons, Net3 has been referred to here as the “Abstract-Knowledge system” of language.

Even if their functional role in cognition is distinct, Net2 and Net3 are both involved in high-level cognition. They display similar network features in terms of hub properties, with a very high proportion of “satellite” key regions compared to other Nets (Table S4, Appendix S2). Satellite centers are regions whose functional communication supports dialogue between components (van den Heuvel & Sporns, 2013). In our case, they favor communication with regions belonging to other Nets, facilitating multimodal integration or information linking over the course of the tasks. Moreover, we have observed specifically in Net2 and Net3 a clear tendency to reconfigure according to the linguistic demand (i.e., versatile networks with a flexible modular configuration that depends on the language subprocesses involved; Figure 4A). This is consistent with studies showing that these systems are rather auxiliary and differentially involved depending on the nature of task (Fedorenko & Thompson-Schill, 2014). For instance, FPN/MDN is functionally active in controlled and challenging semantic tasks but not in less demanding linguistic tasks (Diachek et al., 2020), which is consistent with the supposed role of FPN in attentional processes and fluid cognition (Assem et al., 2020; Perrone-Bertolotti et al., 2020). In the same line, the Net3 configuration is more likely to be engaged primarily in tasks involving the projection of spontaneous and self-oriented thoughts (such as in verbal mind wandering; Andrews-Hanna, Smallwood, et Spreng 2014; Binder et Desai 2011; Humphreys et Lambon Ralph 2015; Konishi et al. 2015; Lau et al. 2013; Raichle 2015; Wang et al. 2020). Interestingly, the brain spatial distribution of Net2 and Net3 specifically corresponds to the mapping of the 5HT2A receptors involved in serotoninergic transmission (Figure 4B; Savli et al. 2012 ; see also Beliveau et al. 2017, for a high resolution and *in vivo* brain atlas of the serotoninergic system), capable of amplifying or sustaining cortical excitation (Puig & Gulledge, 2011). These receptors indeed modulate whole-brain connectivity, promote flexibility between brain states and processes (Jancke et al. 2021), and thus constitute a relevant biomarker of the functional flexibility as evidenced in Nets 2-3.

However, the two networks are distinct, underpinned largely by specific anatomic connectivity (Figure 4C). In addition, they are differentially sensitive to gender (Figure 4D). Hubs of Net3 show higher FC during tasks in females compared to males, which is consistent with whole-brain FC studies showing that the “females-greater-than-males” regions are mostly located within the DMN (Liang et al., 2021). It is also in line with studies highlighting gender differences on language tasks. Females rely more on a supramodal language network during linguistic processing, whereas men tend to process information in modality-specific cortical regions (Burman et al., 2008; Kaushanskaya et al., 2011). Finally, Net2 and 3 are both subject to the pressures of age but here again differently (Figure 4C). Net2 is negatively impacted by aging. The older the age, the less Net2 regions are functionally connected. This reduction with age in attentional-controlled FC is consistent with the alteration in executive functioning traditionally observed in older subjects (Reuter-Lorenz et al., 2016). Net3, on the contrary, has a high share of hubs that are more densely connected with age, which may reflect a compensatory pathway traditionally observed in aging concerning language (i.e., a semantic strategy: Baciu et al. 2021).

The last system, Net4, holds bilateral sensorimotor cortico-subcortical brain areas. This fourth component of the language connectome is distinct from the perceptual and motor auditory-verbal structures included in Net1 (Figure 2B) but could be an important part of the action-perception circuits of language. The brain regions involved in Net4 have already been described as engaged in several sensorimotor aspects related to language production, in particular: general action selection (premotor); motor execution (SMA); orofacial motor activity (precentral and postcentral language areas); or even timing of motor outputs (putamen and cerebellum; Price 2012 for an exhaustive overview). Besides the primary and secondary sensorimotor regions, Net4 also encompasses a large part of the precuneus. Precuneus supports a high level of interconnectivity with other brain regions, which has led to the identification of functional subdivisions (posterior-visual; central-cognitive/associative; anterior-sensorimotor; Margulies et al. 2009) and has indeed been considered as an important sensorimotor connector hub of the language connectome in our analysis (Table S4, Appendix S2). Importantly, it is a crucial site of production-comprehension coupling in natural speech (Silbert et al., 2014). In addition to speech production, Net4 can indeed be engaged in language comprehension. Semantic grounding, i.e., the semantic links between words and their actions, referent objects and related concepts, appears to depend on semantic circuits that bring together both the circuits related to word form (perisylvian, Net1) and conceptual circuits that underlie, among other aspects, sensory-motor experience (extrasylvian, including Net4). The involvement of the motor system in speech perception and understanding has been observed in various contexts (Fernandino et al. 2022; Schomers et Pulvermüller 2016; Skipper, Devlin, et Lametti 2017 and see Pulvermüller 2018 for the hypothesis of neural reuse of action perception circuits in language), which may explain why Net4 is globally engaged regardless of the subprocess involved in the language tasks (Figure 4A). Finally, several neurotransmitters are involved in the regulation of the activity of sensorimotor regions. However, we observed a specific spatial matching between the sensorimotor language system and the noradrenergic receptor mapping (Figure 4B). Catecholamine noradrenaline has indeed substantial projections to somatosensory and motor areas including primary cortices and the modulatory effects of noradrenaline on sensorimotor processing are diverse. If its contribution to the modulation of arousal states (Holland et al., 2021) and in adapting sensory circuits for optimal behavior in animals is well documented (see Jacob et Nienborg 2018 for a review), its precise function in humans and in language remains to be investigated.

Overall, our observations reinforce and complement past observations about the neurocognitive architecture of language. The concept of multiple language networks (Hagoort, 2019) or the “theoretical” subdivision of the vast language network into a key system accompanied by several additional systems (or margins; Hertrich et al. 2020) have been previously discussed. It is interesting to note that even if the number of proposed networks varies according to methods used or to primary theoretical frameworks, the task-based networks or systems defined herein are broadly consistent with previous partitioning proposals. The great benefit of a partitioning emerging directly from data is to pinpoint latent mechanisms that transcend our classical cognitive descriptions (see interesting discussions on the current problem of brain-behavior concordance or the blurriness and ambiguity associated with terminology and definition of psychological constructs: Anderson 2011; Buzsáki 2020). Data-driven ontology provide an independent view, here from a neuro-centric perspective (Roger et al., 2022) and can serve as “*lingua franca across disciplines and theoretical gaps*” (Eisenberg et al., 2019).

However, since they are derived from the observations, these partitions depend directly on the quantity, quality, sensitivity and validity of the data used. This study has the advantage of being based on a database including a rich diversity of fMRI protocols, varying on a wide range of language characteristics (Figure 1). However, task paradigms are less easy to implement than the resting state. They are often very controlled, require multiple repetitions to obtain a robust signal/noise ratio, are more prone to movement artifacts, and induce higher interindividual variability (Park et al., 2020). This often leads to smaller final samples, which may be an important issue for subsequent analyses. Taking into consideration the need to maximize observations to ensure robust results, we have focused most of our analysis on the investigation of the general connectome or on the subprocesses common to several tasks (and not on individual tasks). The compilation of even larger databases will allow for broader and more detailed investigations. For example, it would be important to take into account the pragmatic aspects of language (Rasgado-Toledo et al., 2021), which are not specifically valued in the *InLang* database. Moreover, the current trend is to extend the framework of fMRI paradigms traditionally employed in “laboratory” settings to less controlled and more ecological protocols (Verga & Kotz, 2019). As evidenced by the recent Neuroimage Special Issue (Finn et al., 2022), several initiatives and datasets are steering towards accounts of cognition in more natural settings (e.g., Bhattasali et al. 2020; LeBel, Jain, et Huth 2021; Nastase, Goldstein, et Hasson 2020 for language). Similarly, (neuroimaging) multimodal initiatives have flourished in recent years [e.g., HCP: Van Essen et al., (2013); UK Biobank: Sudlow et al., (2015); ENIGMA: Thompson et al., (2020) ; CamCAN : Taylor et al., (2017) data collections]. In this direction, multimodal datasets including language tasks performed in both fMRI and MEG for example are interesting to address more directly the neurobiological correlates of language, in terms of anatomy or temporal evolution (i.e., dynamics; e.g., the MOUS dataset: Schoffelen et al. 2019). However, the (language) tasks included in such multimodal datasets are still generally limited.

The present study offers a language atlas that relies on a thorough topological (i.e., spatial) analysis of FC. The next step is to identify the causal organization (e.g., hierarchies/heterarchies) and precise timeframes in which its components/regions engage depending on subprocesses/mechanisms at work. MEG or electroencephalography (EEG; placed on the scalp, the cortex or intracranial) are valuable tools for the evaluation of such dynamics. To this end, and using intracranial EEG recordings, a recent study examined the dynamic organization of naming (Forseth et al., 2021). They observed that regions were mainly co-activated during extended periods, confirming that complex behaviors such as speech production requires the coordination of discrete network states (defined as set of reference dynamics that coordinate the generation and transmission of information throughout the cortex; see also the concept of meta-networking of Herbet & Duffau, 2020). They were able to sequence, map and identify the temporality of the different transient states; globally confirming the seminal model of word production proposed by Indefrey & Levelt, 2004). Ultimately, the definition of a comprehensive repertoire of language states and causal relationships (i.e., effective and directed functional connectivity; e.g., Deco, Vidaurre, et Kringelbach 2021) in various tasks may extend our understanding of language functioning. Finally, although we explored a number of modulators, a study of LANG properties on data acquired in multilingual individuals (Li et al., 2021), in pediatric populations (Wang et al., 2022), or in pathology in relation with neuropsychology (neuroplasticity and neurocognitive efficiency; e.g., Banjac et al. 2021) would also enrich our findings.

## 4 Conclusion

Language is a multi-faceted cognitive function. In an attempt to account for the multidimensionality of language, we performed functional connectivity analyses on a multi-paradigm fMRI database (*InLang*), gathering thirteen different language tasks. It allowed us to inspect the language connectome in depth, in particular on its spatial properties and functional attributes. In all, this study reaffirms that high-level cognition such as language emanates from synergistic exchanges of external and internal information across specialized systems. Our results highlight the involvement of essential discrete networks (or components) that are settled around a core “language-related” system. In addition, the flexible engagement of some key regions depending on several modulating factors such as the linguistic demand points to the dynamic nature of language. From ontology-based data integration, we propose a connectomic atlas of the “language mosaic”, which can serve as a reference for investigating additional conditions or pathologies altering language functioning.

## 5 Materials and Methods

### 5.1. Dataset

The *InLang* database contains data of 13 different linguistic tasks from 7 previously published fMRI protocols (Baciu et al., 2016; Banjac et al., 2020; Grandchamp et al., 2019; Haldin et al., 2018; Hoyau, 2018; Perrone-Bertolotti et al., 2011, 2015, 2017; Perrone-Bertolotti et al., 2012), as well as respective structural MRIs (T1w). In all, 359 functional scans have been acquired between 2010 and 2019 from 150 healthy adults (all right-handed; 64 females: F/86 males: M; Table S1 contains the subjects’ characteristics, by tasks). The database includes 114 “young” (18-59; 50F/64M) and 36 “senior” (60-85; 14F/22M) volunteers (Figure 1). fMRI and T1w scans of all participants have been formatted in BIDS standard and preprocessed using conventional tools (Appendix S1).

### 5.2. Task-based connectomes

Regions of interest (ROIs) covering both the brain and the cerebellum were defined from 6 mm radius spherical regions built around the 264 coordinates in MNI space proposed by Power et al. (2011). The images used for signal extraction (beta values) were the statistical parametric maps containing the linear contrasts between the HRF parameter estimates for the conditions of interest (henceforth “contrast images”). Nilearn (https://nilearn.github.io/stable/index.html; Abraham et al., 2014) was used to delineate ROIs and extract the signal. Mean signal for each of these ROIs was extracted by participant and task for each project separately (Appendix S1).

We then estimated task-based connectomes for each task individually by correlating the beta signals extracted from all nodes (264 ROIs; Pearson correlations). These matrices of functional connectivity (FC) were thresholded. We applied a 5% threshold, which defines the 5% of the highest positive correlation values, considered to represent non-spurious internodal connections. The matrices were binarized: 1 was assigned to internode connections which survived to the given density threshold, and 0 was assigned otherwise. Task-based connectomes were thus built from these binarized matrices, reduced to a fixed number of edges (top 5%, 3485 edges). Different graph metrics were computed on the task-based connectomes: global (network-wide), intermediate (modularity) and local (nodal), using Networkx (https://networkx.org/).

### 5.3. Network measures

#### 5.3.1. Global connectivity profiles

To determine the global functional connectivity (FC) profile of the language tasks, we computed several parameters at the global level (i.e., network-wide estimates), namely:

- The global efficiency (E_glob_) as proposed by Latora & Marchiori (2001), which is the average of the unweighted efficiencies over all pairs of nodes:

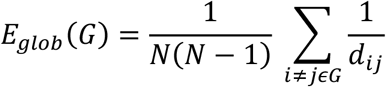

where *N* is the total number of nodes in the network *G*, the distance d(*i,j*) corresponds to the number of edges in a shortest path between any two nodes *i* and *j*. E_glob_ represents the capacity of a given network to efficiently integrate and transmit information between the network components or subnetworks (e.g., Bullmore et Sporns 2012; Roger et al. 2019; Stanley et al. 2015). The higher the value, the more likely that information transfer is fast.
- The local efficiency (E_loc_) which is the average of the local efficiencies of each node. Local efficiency of a node *i* corresponds to the average global efficiency of a subgraph induced by the neighbors of *i* (Latora & Marchiori, 2001):

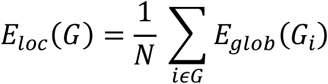

E_loc_ reveals the network’s tendency to effectively share information within immediate local communities or the capacity of a given network to segregate the information processing (e.g., Bullmore et Sporns 2012; Roger et al. 2019; Stanley et al. 2015). The higher the value, the more locally efficient the network is.
- The integration–segregation balance (I:S) as expressed by the difference between E_glob_ and E_loc_ (E_glob_ – E_loc_). The integration–segregation balance allows to estimate how the functional organization of a task promotes either (1) more independent processing of specialized subsystems (i.e., segregation) or (2) cooperation between different subsystems (i.e., integration; Wang & al., 2021). A positive balance reflects a network with a general tendency toward functional integration (E_glob_ > E_loc_) while a negative balance reflects a general tendency toward functional segregation (E_loc_ > E_glob_).
- The mean geodesic cortical distance (*d̄*) between functionally interconnected nodes (*d̄*(G)). We extracted the relative spatial layout of regions along the cortical surface by using existing scripts (https://github.com/margulies/topography/tree/master/utils) based on an algorithm developed to approximate the exact geodesic distance from triangular meshes (Oligschläger et al., 2017). “Physical” geodesic distances between pairs of nodes, estimated in mm, were quantified from the Power’s nodes coordinates, projected onto a template surface mesh (fsaverage5). This resulted in a node-by-node matrix of geodesic cortical distance. From the geodesic distance matrix, we averaged the distribution of distance-to-connected-areas of the relevant functional connections identified from the thresholded FC matrices. We thus obtained the global geodesic distance of the functional connectivity for each of the networks corresponding to the different language tasks. The higher the mean geodesic distance between functionally connected regions, the further apart the connected regions are on average along the cortical surface.

#### 5.3.2. General LANG connectome and subprocesses

We have estimated the similarity of the global FC profiles of the different language tasks based on the Euclidean distance. We applied data-driven hierarchical clustering approach on the similarity matrix and estimated the partitioning. Thus, we identified categories of tasks with more or less similar connectivity profiles or global network topology. The internal composition and nature of the tasks assigned to each of the identified groups can reveal putative linguistic subprocesses that may be latent and common to several tasks (see also Appendix S1 for a rationale and a similar method applied to functional activation maps). Starting from the partitioning, we estimated the FC matrices of each of these main task groups – hereafter referred to as subprocesses – from the scans of the respectively involved tasks and using the same procedure described above (Section 5.2.).

In addition, and still following the same procedure, we generated the general task-based language connectome – abbreviated LANG – corresponding to the FC matrix derived from all language tasks (and thus also from the subprocesses). Only scans corresponding to the “younger” *InLang* cohort were considered for the calculation of task, subprocess, and LANG FC matrices. Scans from the “older” group were used for other analyses described later. The global FC parameters reported in the previous section (Section 5.3.1) were also extracted for LANG and the subprocesses.

#### 5.3.3. Intermediate composition of LANG

We performed modularity analyses to determining community structure of the general LANG connectome. We used the Louvain community detection algorithm (Blondel et al., 2008) implemented in Networkx (https://networkx.org/). Louvain’s method is widely used for community detection in neuroscience and has been previously shown to outperform other community detection methods (Yang et al., 2016). To ensure the stability of the final partition, we repeated the modular partitioning process 100 times (Schedlbauer & Ekstrom, 2019) and we evaluated the best LANG partition (Aynaud, 2018) on the matrix averaging the results of all iterations. Each ROI was assigned to a specific community (i.e., a subnet, here denoted LANG “Net”). To facilitate subsequent analyses and interpretations, the Power’s coordinates of LANG ROIs were mapped to the HCP’s multimodal parcellation (version 1.0: HCP_MMP1.0 proposed by Glasser et al. 2016), which consists of 180 brain parcels. Cerebellum coordinates were mapped to the probabilistic human cerebellum atlas SUIT (Diedrichsen et al., 2009).

Still concerning intermediate configurations between the global level of the network and the nodal ROIs level, we focused on identifying densely interconnected subgraphs of LANG. Complete subgraphs, called cliques, are all-to-all connected sets of brain regions providing architecture that isolates information transmission processes (Giusti et al., 2016) and supports efficient and specialized processing (Sizemore et al., 2018). A maximal clique is one that includes the largest possible number of nodes and to which no more nodes can be added. Using Networkx, we estimated ω(G) which is the number of nodes in a maximal clique of G and marked the relevant nodes.

#### 5.3.4. Nodal properties of LANG

We calculated the degree centrality (DC; denoted *k*) of each node *i* as the number of adjacent edges to the node (*ki*), from the reduced and unweighted adjacency LANG matrix. DCs are a convenient metric for highlighting brain regions with a high degree of connectivity or “hubs”, and form the basis for other measures of nodal graph theory.

From the DCs (k), we computed the within-component degree z-score (*zi*), that expresses the extent to which node *i* is connected to other nodes in its respective component and is calculated as follows (Guimerà & Nunes Amaral, 2005):

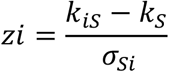

with *k_iS_* the number of connections of node *i* to the other nodes in the subgraph component *S* (i.e., the Net) and *k_S_* and σ*_Si_* respectively the mean and SD of the within-component DC over all nodes in *S*.

To quantify to what extent a node connects across all components, we measure the participation coefficient (*PCi*). The following conventional formula (Guimerà & Nunes Amaral, 2005) was applied, with *m* the set of components *S* or Nets (here 4):

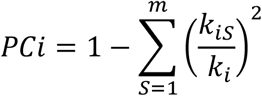

Following Schedlbauer & Ekstrom (2019) and because of the narrow distribution of the *PC*s we z-scored the coefficients (zPCi) from each network.

The *zi* and *zPCi* values have enabled to assign a specific role to each of the LANG ROIs. The nodes were classified according to their type of functional communication within the connectome as follows: connector (high *z_i_*/high z*PC_i_*; high intra-Net and high inter-Net FC); provincial (high *z_i_*/low z*PC_i_*; high intra-Net FC); satellite (high *z_i_*/low z*PC_i_*; high inter-Net FC); or peripheral (low *zi*/low *zPCi*; low inter-Net FC). We applied this classification as proposed in previous studies (Bertolero et al., 2015; Cohen & D’Esposito, 2016; van den Heuvel & Sporns, 2013) and with *z_i_* > 0 corresponding to “high *z_i_*” and z*PC_i_* > 0 corresponding to “high *P_i_*”.

Finally, we were interested in quantifying the rich club organization of the networks. A rich club reflects a set of nodes in the network of whose level of interconnectivity (i.e., richness) exceeds the level of FC that can be expected by chance. For each degree k, the rich-club coefficient (*ϕ*) is the ratio of the number of actual to the number of potential edges for nodes with degree greater than k (Colizza et al., 2006):

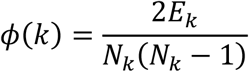

where *Nk* is the number of nodes with degree larger than *k*, and *Ek* is the number of edges among those nodes.

We compared and normalized the rich club coefficient to sets of “equivalent” random networks. An empirical null distribution constituted from the average of 1000 random networks of equal size and degree distribution was generated (*ϕrand* (*k*)).

The difference between *ϕ*(*k*) and *ϕrand*(*k*) allowed us to obtain the normalized rich club coefficient *ϕnorm*(*k*):

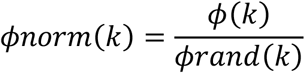

In line with previous work (Colizza et al., 2006; Grayson et al., 2014; Heuvel & Sporns, 2011), a network was considered to have rich club organization when *ϕnorm* was greater than 1 for a continuous range of increasing k (rich club regime). Rich club nodes were brain regions taking part in these densely connected networks (or rich clubs), forming a functional unit. We considered as rich club hubs the nodes taking part in the club at value k where the strongest rich club effect was observed.

### 5.4. Statistics

#### 5.4.1. Hemispheric asymmetry

The FC hemispheric asymmetry of the ROIs was estimated with the DCs. We derived a connectivity-based lateralization index (LI), by contrasting the k values of homotopic nodes (comparison of FC between mirror areas), according to the following formula:

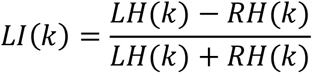

With LH(k) being the DC for the ROI in left hemisphere, RH(k) the DC for the homolateral ROI in the right hemisphere.

We also calculated global LIs at the connectome or Net level by averaging the corresponding nodal LIs. LI values can range continuously from −1 to 1 and the following landmarks were considered for interpretation: −1 = complete RH dominance; +1 = complete LH dominance and between −0.2 and +0.2 = no clear dominance (Roger, Pichat, Renard, et al., 2019; Rolinski et al., 2020; Seghier, 2008).

#### 5.4.2. Functional and structural matching

We used mappings provided by previously published tools to estimate the spatial concordance between LANG and (1) the neurotransmitter pathways; or (2) the terminations of large white matter (WM) bundles. The “functional” maps are issued from nuclear imaging-derived neurotransmitter maps implemented in the JuSpace toolkit (Dukart et al., 2021), specifically designed to link neuroimaging (MRI data) with underlying neurotransmitter information (as revealed by PET and SPECT tracers). The “structural” maps came from the deep-learning algorithm TractSeg (Wasserthal et al., 2018), which offers the segmentation of the main long-range WM brain bundles. It also allows the generation of grey matter masks that are linked by the bundles (ending masks). These ending masks were used here to define the structural connection maps of each bundle or combination of bundles.

The functional and structural maps were registered to the surface template and binarized. Each parcel of the HCP_MMP1.0 template was coded according to the presence/absence of map coverage, with: 1 corresponding to at least 40% coverage of the parcel surface (> 40%); and 0 to less than 40% coverage (< 40%).

We then used the simple matching coefficient (SMC; Boriah et al., 2008; Sokal & Michener, 1958) method to quantify the spatial concordance between LANG and each of the functional and structural binary maps. SMC indicates the coincidence ratio between the mutual presences (and absences) and the length of the binary sequences: 0% means that the labels have nothing in common and 100% that they have identical sequences. Only coefficients exceeding 2/3 of the total agreement (SMC > 0.67) were considered relevant.

#### 5.4.3. Cross-processes flexibility

We computed a flexibility index by using multilayer network model and with a method close to that of Betzel et al. (2017). The layers of the model were constituted from the matrices of the 5 groups of tasks (i.e., subprocesses) identified with data-driven clustering analyses (see Section 5.3.2). To keep a common reference across the layers, the matrices were restricted to the LANG 131 ROIs and re-estimated on this basis. We applied the generalized Louvain package (Jeub et al., 2011), suited to determine community structure in multiplex graphs (Bassett et al., 2011, 2013; Mucha et al., 2010). This method has the advantage of preserving the community labels consistently across layers (here the task groups), avoiding thus the issue of community matching (Yang et al., 2021).

From the communities assigned across layers, we calculated a flexibility score as previously proposed by (Bassett et al., 2011). Flexibility f_i_ of a node corresponds to the number of times that a node changes its modular assignment across layers, normalized by the total number of possible changes (i.e., the total number of layers minus 1, here 4). In short, the f-score reflects the frequency a brain region changes its community assignment. It ranges from 0 to 1, where 0 corresponds to a region that never changes module whatever the subprocess/task involved (stable across all layers); and 1 corresponds to a region that never belongs to the same module on the 5 layers. We also calculated the mean flexibility (F) over all nodes in the network to examine the global flexibility of the system.

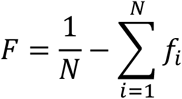

#### 5.4.4. Inter-individual variability

We assessed inter-subject variability by considering the individual signal values, extracted for each (young) individual and on each of the LANG ROIs. In order to remove the variability induced by the task, we normalized the beta values, considering the mean and standard deviation of the other subjects who performed the given task (z-score betas). When individuals performed multiple tasks (for subjects enrolled in the same protocol), we averaged the z score betas for these subjects to avoid accounting for additional intra-individual variability. We thus performed the measurement of inter-individual variability by considering the subjects and not the scans. In addition, we considered only the “young” participant cohort to limit the age effect. The average and absolute z scores of each region were then divided into 3 bins of increasing interindividual variability.

#### 5.4.5. Age effect

To highlight LANG ROIs that are the most resilient/vulnerable to the aging effect, we examined the subjects from our two “young” and “old” cohorts who performed the same task (NAM: NAM young; NAM old). Age was considered continuously in our statistical analyses, from age 20 to 85 (n = 82). We computed standard correlation coefficients (Pearson *r*) between the age and DC (here estimated on the basis of individual connectivity matrices). This allowed us to observe a positive (positive and high *r*) or negative (negative and high *r*) age effect on task-based FC.

#### 5.4.6. Gender effect

We estimated the gender effect by generating the FC matrices reduced to the LANG ROIs for males (M) and females (F; self-reported gender) separately. The average DCs obtained for each ROI were then compared between the two groups. Only the top 10% of the largest differences (Males>Females and Females>Males, independently) were retained to define the most diverging ROIs.

## 6 Supporting information (SI)

**Appendix S1: Material and Methods SI**

Description of the InLang database

fMRI protocol details

Participants information

Data acquisitions & (pre)processing

**Appendix S2: Results SI**

Supplementary tables (global and nodal properties)

LANG package files

LANG maps (Nets, correlates, variability)

## Acknowledgements

We thank Philippe Kahane, Lorella Minotti, Olivier David, Arnaud Attyé, Laurent Lamalle and Jean Francois Démonet for their support and contribution to the NeuroCoG project. We also thank Cédric Pichat, Sonja Banjac, Laurent Torlay and Eric Guinet for their help in data collection and their useful insights.

