## Appendix S1 for "Unraveling the functional attributes of the language connectome: crucial subnetworks, flexibility and variability"

#### Supplementary information (SI) – Materials and methods

#### *InLang* database, fMRI paradigms and functional activation maps

This study and the *InLang* database include data from one hundred and fifty healthy adult participants of seven previously published research projects, which are: NEREC (Perrone-Bertolotti et al., 2015), NeuroMod (Hoyau, 2018), Recovery (Haldin et al., 2018), Plast-LANG (Perrone-Bertolotti et al., 2011, 2013, 2017), Reorg-GEREC (Banjac et al., in press), SEMVIE (Baciu et al., 2015) and InnerSpeech (Grandchamp et al., 2019). Materials and methods will be separately described by project whenever those differ.

### Participants

All participants included in this study were native French speakers with normal or corrected-to-normal vision and no history of neurological or psychiatric disorders. Their characterization by sample size, age and gender by project is as follows: NEREC (n = 13, mean age: 23.6 ± 2.96 years, 11 females), NeuroMod (n = 28, mean age: 66.57 ± 7.39 years, 15 females), Recovery (n = 11, mean age: 33.9 ± 11.31 years, 5 females), Plast-LANG (n = 24, mean age: 27.2 ± 3.7 years, 12 females), Reorg-GEREC (n = 20, mean age: 21.35 ± 3 years, 9 females), SEMVIE (n = 30, mean age: 56.1 ± 17 years, 9 females) and InnerSpeech (n = 24, mean age: 28.4 ± 10 years, 13 females). Table S2 below provides the main characteristics of each subject, by project. Participants were predominantly right-handed according to the Edinburgh Handedness Inventory (Oldfield, 1971).

The studies were approved by the local Ethics Committee "Comité de Protection des Personnes" with the following approval codes: NEREC (CPP: 09-CHUG-14, 04/06/2009), NeuroMod (CPP n° 2017-A00834-49), Recovery (CPP-ISIS 07PHR04-N°DCIC/06/25), Plast-LANG (CPP: 09-CHUG-14, 04/06/2009), Reorg-GEREC (CPP 09-CHUG-14; MS -14-102), SEMVIE (CPP: 2014-A00569-38). The InnerSpeech project was approved by the local Ethics Committee "Comité de Protection des Personnes" Sud-Est V and by the National Competent Authority France-ANSM (Ref. CPP: 14-CHUG-39, Ref. Promoteur: 38RC14.304, ID-RCB: 2014-A01403-44, Ref. ANSM: 141200B-31, ClinicalTrials.gov ID: NCT02830100) (CPP: 09-CHUG-14, 04/06/2009). All participants gave informed consent before volunteering.

### Functional MRI paradigm

##### *Stimuli and tasks*

###### NEREC

- **In the object naming task (NAM)**, participants were instructed to overtly name 80 black and white drawings of objects and animals from the DO80 database (Metz-Lutz et al., 1991).

The run consisted of four task periods and four control periods of 85 s, with a total duration of 11 minutes and 20 seconds. Each task period contained 20 stimuli and 10 fixation crosses of

2.5 s. Control periods were similarly built, but the stimuli were grey square or round geometrical shapes instead, to which participants were instructed to simply say “square” or “round”.

In this study, we contrasted NAM to the baseline to consider all NAM tasks together. When contrasted with the **baseline**, the NAM task involves **word production** and engages **semantic** processes.

#### NeuroMod

- **In the object naming task (NAM)**, participants were instructed to overtly name 48 black and white drawings of objects and animals from the DO80 database (Metz-Lutz et al., 1991).

The run consisted of four task periods and four control periods of 60 s, with a total duration of 8 minutes. Each task period contained 12 stimuli, and each trial consisted of the presentation of a fixation cross for 0.5 s and a stimulus for 4.5 s. Control periods were similarly built, but the stimuli were scrambled versions of the images used for the task periods, to which participants were instructed to simply say “bababa”. Two sessions with two runs each were carried out, with one of the sessions for the condition of anodal tDCS and the other one for the condition of sham, in counterbalanced order across participants. For the current study, we only included data from the sham session and from the first run of the anodal tDCS session.

In this study, we contrasted NAM to the baseline to consider all NAM tasks together. When contrasted with **the baseline**, the NAM task involves **word production** and engages **semantic** processes.

#### Recovery

Participants performed three language tasks: object naming (NAM), word repetition (REP) and rhyme detection (RHYM); each in a separate run. Each run had a block design and alternated task and control periods of 24 s which contained 4 stimuli. One fixation cross of 1 s was also presented for each block and 2 fixation crosses of 3 s between blocks. Oral responses were recorded with an MRI compatible optical microphone (FOMRITM III, version 1.2).

- **For the object naming task (NAM)**, participants were instructed to overtly name black and white drawings of objects and animals from the DO80 database (Metz-Lutz et al., 1991).

Control stimuli were grey square or round geometrical shapes, to which participants were instructed to simply say “square” or “round”.

In this study, we contrasted NAM to the baseline to consider all NAM tasks together. When contrasted with **the baseline**, the NAM task involves **word production** and engages **semantic** processes.

- **For the repetition task (REP)**, participants were instructed to repeat French mono-syllabic words which consisted of the 10 French oral vowels /i, e, ɛ, a, y, ø, oe, u, o, ɔ/ (such as “où”

/u/, *where*) and of 10 obstruent consonants /t, d, s, z, ʃ, ʒ, k, g, l, ʁ/ followed by the vowel /a/ (such as “gars” /ga/, *guy*).

These words were pronounced by a female voice and transmitted to participants via MRI-compatible headphones. White noise was presented during the control blocks, with no response required from participants.

When contrasted with the **control task**, the REP task involves **overt word production** and engages **semantic** processes.

- **For the rhyme detection task (RHYM)**, participants were instructed to judge if pairs of words rhymed or not by pressing the corresponding button.

Half of these pairs was phonologically and orthographically congruent, and the other half was phonologically and orthographically incongruent (Baudo & Vernisse, 2001). Words were presented in black font and placed one above the other. During control blocks, two horizontal lines were presented, with no response required from participants.

When contrasted with the **control task**, the RHYM task involves **phonology decoding** and engages **monitoring** processes.

#### Plast-LANG

Participants performed three language tasks: phoneme detection (PHON), semantic categorization (SEM) and prosody judgment (PROS), each in a separate run. All tasks had an event related design. Runs contained target and control stimuli, with 30 items of each type for the first two tasks and 48 items of each type in the prosody judgment task. The inter-trial interval in the phoneme detection and in the semantic categorisation task had 2.5 s, corresponding to a 0.5 s fixation followed by a 2 s stimulus. The inter-trial interval in the prosody judgment task had on average 4 s. Additional null events consisting of a fixation cross were included in all runs, being 35 for the first two tasks and 30 for the last one. This gave a total run duration of approximately 9 min in all cases.

- **For the phoneme detection task (PHON)**, participants were instructed to judge if the written pseudo-word presented contained a sound /o/ by pressing the corresponding button for “yes” or “no”.

- **For the semantic categorisation task (SEM)**, participants were instructed to judge if the written word presented belonged to a living or a non-living category by pressing the corresponding button. To perform these tasks, participants were required to pronounce the items silently and without articulation.

For both tasks, the control condition consisted of a low-level visual detection task. Participants should indicate whether the unreadable words presented contained any character whose font size was larger than the others in the same word.

When contrasted with the **control task**, the PHON task involves **phonology decoding** and engages **monitoring** processes.

When contrasted with the **control task**, the SEM task involves **word comprehension** and engages **semantic** processes.

- **For the prosody judgment (PROS)**, participants had to judge if a sentence had narrow focus or if it was neutral by pressing the corresponding button.

This aim was reached via an indirect request which consisted in asking participants to judge if the speaker uttered the sentence to convey a correction by stressing part of it, or if the speaker uttered the sentence in a neutral fashion. Forty-eight target sentences had narrow focus, being half of them with focus on the subject and the other half with focus on the object. Control sentences were uttered in a neutral fashion. All sentences were transmitted to participants via MRI-compatible headphones.

When contrasted with the **control task**, the PROS task involves **prosody decoding** and engages **monitoring** processes.

#### Reorg-GEREC

- **In the sentence generation task (GENE)**, participants were instructed to covertly generate a sentence for each of the words they heard through MRI-compatible headphones.

They were required to keep repeating the sentence generated until they heard the next word. Forty black and white drawings from the DO80 database (Metz-Lutz et al., 1991) were used. The run consisted of five task blocks and five control blocks separated by five “rest” blocks, with a total duration of 7.3 minutes. Each task block contained eight stimuli and the inter-trial interval was of 5 s. Control blocks were similarly built, but a pseudo-word was repeated 8 times instead, and participants were instructed to listen and to try not to talk covertly. Five “rest” blocks were also included in the run and consisted of the presentation of a fixation cross for 10 s. The “rest” blocks were placed after each task block, and the order of blocks within the run was therefore task, rest and control.

When contrasted with the **control task**, the GENE task involves **lexico-syntactic production** and engages **semantic** processes.

#### SEMVIE

Two language tasks, object naming (NAM) and verbal fluency (FLU), were performed in separate runs with a block design. Oral responses were recorded with an MRI compatible optical microphone.

- **For the object naming task (NAM)**, participants were instructed to overtly name 80 black and white drawings of objects and animals from the DO80 database (Metz-Lutz et al., 1991).

The run alternated four task periods and four control periods of 50 s, separated by a 0.5 s fixation cross, and had a total duration of 7 minutes and 6 seconds. Each task period contained 20 stimuli. Control periods were similarly built, but the stimuli were grey square or round geometrical shapes instead, to which participants were instructed to simply say “square” or “round”.

In this study, we contrasted NAM to the baseline to consider all NAM tasks together. When contrasted with **the baseline**, the NAM task involves **word production** and engages **semantic** processes.

- **For the verbal fluency task (FLU)**, participants were instructed to overtly generate as many words as possible belonging to the proposed semantic category.

The semantic category (animals, clothes, vegetables, and sports) was presented visually as a written word at the beginning of each block, which lasted 1 minute. The task blocks were alternated with control blocks of 30 s during which participants were required to fixate a cross and to try not to generate any words. The run lasted 6 minutes.

When contrasted with the **control task**, the FLU task involves **word production** and engages **semantic** processes.

#### InnerSpeech

Participants performed a sequence of five language tasks: Speech Perception (SP), Monologal Self-voice (MS), Monologal Other-voice (MO), Dialogal Other-voice (DO) and Verbal Mind Wandering (VMW), which was repeated in the same run for three times. Participants performed two runs with this design, and the order of tasks in the sequence was counterbalanced across participants. Each run had a block design with three sequences of five task blocks. Four blocks (SP, MS, MO and DO) contained 5 stimuli each, with an inter-trial interval of 6 s and stimuli duration of 2 s. The VMW block started with a similar 2 s stimulus and lasted 32 s. One fixation cross of 2 s was presented between trials within blocks, and another one of 8 s was presented between blocks.

Participants were first introduced to a high-pitched voice avatar, who provided the instructions for the tasks. For the speech perception (SP) task and the intentional inner speech tasks (MS, MO and DO), each trial started with a 2 s stimulus presentation of a written word together with its pictorial illustration. The picture was framed within a clock, which rotated during the 4 s following stimulus presentation during the period of participant’s response.

- In the **speech perception (SP) task**, participants had to listen to the word’s definitions provided by the avatar through MRI-compatible headphones.

- In the **monologal self-voice (MS) task**, participants had to covertly generate their own definitions.

- In the **monologal other-voice (MO) task** the instruction was similar to that in the ISS task, but participants should imitate the avatar’s voice.

- In the **dialogal other-voice (DO) task**, participants had to imagine that the avatar was addressing them and naming each object to them. Four lists of 30 nouns were created with the LEXIQUE database (New et al., 2001) for the tasks.

- In the **Verbal Mind Wandering (VMW) task**, the block started with a 2 s stimulus similar to those presented in the other tasks, but the instruction for the remaining 30 s was that the participants should fixate the rotating clock and monitor the occurrence of spontaneous verbal thoughts by indicating their time on the clock with a joystick.

When contrasted with the **baseline**, the MS, MO and DO tasks involve **speech control** and engage **monitoring** processes.

When contrasted with the **baseline**, the VMW tasks involves **wandering** states and engage **unintentional** speech processing.

**Table S1:** Summary of the *InLang* projects, participants, tasks, statistical contrasts and main language (sub)functions theoretically targeted (*InLang* database)

| fMRI Projects | Participants (N) | Tasks | Statistical Contrasts | Targeted language subprocesses (theoretical) | | | | |
|---|---|---|---|---|---|---|---|---|
| | | | | **Semantic** Conceptual knowledge | **Decoding** Phonology Prosody | **Production** Lexical Formulation | **Monitoring** Voice & Sound | **Wandering** Spontaneous Thoughts |
| **Plast-LANG** | 24 | Semantic categorization (SEM) | Task *versus* Control | 1 | | | | |
| | | Phoneme detection (PHON) | Task *versus* Control | | 1 | | 2 | |
| | | Prosody detection (PROS) | Task *versus* Control | | 1 | | 2 | |
| **SEMVIE** | 30 | Overt object naming (NAM) | Task *versus* Baseline | 2 | | 1 | | |
| | | Overt categorical fluency (FLU) | Task *versus* Control | 2 | | 1 | | |
| **NeuroMod** | 28 | Overt object naming (NAM) | Task *versus* Baseline | 2 | | 1 | | |
| **NEREC** | 13 | Overt object naming (NAM) | Task *versus* Baseline | 2 | | 1 | | |
| **Reorg-GEREC** | 20 | Covert sentence generation (GENE) | Task *versus* Control | 2 | | 1 | | |
| **Recovery** | 11 | Overt monosyllabic word repetition (REP) | Task *versus* Control | 2 | | 1 | | |
| | | Overt object naming (NAM) | Task *versus* Baseline | 2 | | 1 | | |
| | | Word rhyme detection (RHYM) | Task *versus* Control | | 1 | | 2 | |
| **InnerSpeech** | 24 | Speech perception (SP) of word's definitions | Task *versus* Baseline | 1 | | | | |
| | | Covert monologal sentence with self voice (MS) | Task *versus* Baseline | 2 | | | 1 | |
| | | Covert monologal sentence with self voice (MO) | Task *versus* Baseline | 2 | | | 1 | |
| | | Covert dialogal sentence with other voice (DO) | Task *versus* Baseline | 2 | | | 1 | |
| | | Verbal mind-wandering (VMW) | Task *versus* Baseline | | | | | 1 |
| **ALL** | 150 | | | | | | | |

| NEREC | | | NeuroMod | | | Recovery | | | Plast-LANG | | | Reorg-GEREC | | | SEMVIE | | | InnerSpeech | | |
|---|---|---|---|---|---|---|---|---|---|---|---|---|---|---|---|---|---|---|---|---|
| **ID** | **Gender** | **Age** | **ID** | **Gender** | **Age** | **ID** | **Gender** | **Age** | **ID** | **Gender** | **Age** | **ID** | **Gender** | **Age** | **ID** | **Gender** | **Age** | **ID** | **Gender** | **Age** |
| **sub-01** | F | 22 | **sub-01** | F | 67 | **sub-01** | F | 23 | **sub-01** | F | 29 | **sub-01** | F | 19 | **sub-01** | M | 59 | **sub-01** | F | 25 |
| **sub-02** | F | 23 | **sub-02** | F | 69 | **sub-02** | F | 31 | **sub-02** | F | 29 | **sub-02** | M | 19 | **sub-02** | M | 33 | **sub-02** | F | 20 |
| **sub-03** | F | 23 | **sub-03** | M | 71 | **sub-03** | M | 36 | **sub-03** | F | 33 | **sub-03** | F | 19 | **sub-03** | M | 67 | **sub-03** | M | 27 |
| **sub-04** | F | 28 | **sub-04** | F | 56 | **sub-04** | F | 23 | **sub-04** | F | 25 | **sub-04** | M | 21 | **sub-04** | M | 65 | **sub-04** | F | 38 |
| **sub-05** | F | 27 | **sub-05** | F | 53 | **sub-05** | M | 22 | **sub-05** | F | 30 | **sub-05** | F | 18 | **sub-05** | M | 30 | **sub-05** | M | 26 |
| **sub-06** | F | 22 | **sub-06** | F | 74 | **sub-06** | M | 26 | **sub-06** | F | 25 | **sub-06** | F | 18 | **sub-06** | F | 39 | **sub-06** | F | 46 |
| **sub-07** | F | 23 | **sub-07** | M | 71 | **sub-07** | F | 50 | **sub-07** | F | 25 | **sub-07** | M | 20 | **sub-07** | M | 34 | **sub-07** | F | 20 |
| **sub-08** | F | 28 | **sub-08** | M | 65 | **sub-08** | F | 45 | **sub-08** | M | 25 | **sub-08** | M | 23 | **sub-08** | F | 63 | **sub-08** | F | 33 |
| **sub-09** | M | 19 | **sub-09** | M | 84 | **sub-09** | M | 26 | **sub-09** | M | 28 | **sub-09** | M | 23 | **sub-09** | F | 73 | **sub-09** | M | 41 |
| **sub-10** | F | 23 | **sub-10** | F | 75 | **sub-10** | M | 38 | **sub-10** | F | 25 | **sub-10** | F | 19 | **sub-10** | F | 48 | **sub-10** | F | 26 |
| **sub-11** | F | 22 | **sub-11** | M | 81 | **sub-11** | M | 53 | **sub-11** | M | 26 | **sub-11** | F | 18 | **sub-11** | M | 69 | **sub-11** | M | 47 |
| **sub-12** | F | 20 | **sub-12** | M | 66 | | | | **sub-12** | M | 29 | **sub-12** | F | 29 | **sub-12** | M | 56 | **sub-12** | M | 20 |
| **sub-13** | M | 27 | **sub-13** | M | 66 | | | | **sub-13** | M | 23 | **sub-13** | M | 23 | **sub-13** | M | 52 | **sub-13** | M | 25 |
| | | | **sub-14** | M | 73 | | | | **sub-14** | F | 27 | **sub-14** | F | 25 | **sub-14** | F | 45 | **sub-14** | M | 22 |
| | | | **sub-15** | M | 69 | | | | **sub-15** | M | 30 | **sub-15** | M | 19 | **sub-15** | F | 30 | **sub-15** | F | 19 |
| | | | **sub-16** | F | 58 | | | | **sub-16** | M | 30 | **sub-16** | M | 20 | **sub-16** | M | 56 | **sub-16** | F | 23 |
| | | | **sub-17** | F | 64 | | | | **sub-17** | M | 34 | **sub-17** | M | 22 | **sub-17** | M | 76 | **sub-17** | F | 19 |
| | | | **sub-18** | F | 63 | | | | **sub-18** | F | 24 | **sub-18** | F | 25 | **sub-18** | M | 78 | **sub-18** | F | 23 |
| | | | **sub-19** | F | 62 | | | | **sub-19** | F | 19 | **sub-19** | M | 24 | **sub-19** | M | 79 | **sub-19** | M | 20 |
| | | | **sub-20** | F | 62 | | | | **sub-20** | M | 33 | **sub-20** | M | 23 | **sub-20** | M | 41 | **sub-20** | F | 27 |
| | | | **sub-21** | F | 70 | | | | **sub-21** | M | 31 | | | | **sub-21** | M | 44 | **sub-21** | F | 22 |
| | | | **sub-22** | F | 70 | | | | **sub-22** | M | 24 | | | | **sub-22** | M | 68 | **sub-22** | M | 20 |
| | | | **sub-23** | M | 67 | | | | **sub-23** | F | 22 | | | | **sub-23** | F | 69 | **sub-23** | M | 44 |
| | | | **sub-24** | M | 65 | | | | **sub-24** | M | 27 | | | | **sub-24** | F | 39 | **sub-24** | M | 49 |
| | | | **sub-25** | M | 71 | | | | | | | | | | **sub-25** | M | 84 | | | |
| | | | **sub-26** | F | 58 | | | | | | | | | | **sub-26** | M | 70 | | | |
| | | | **sub-27** | M | 55 | | | | | | | | | | **sub-27** | M | 77 | | | |
| | | | **sub-28** | F | 59 | | | | | | | | | | **sub-28** | F | 69 | | | |
| | | | | | | | | | | | | | | | **sub-29** | M | 36 | | | |
| | | | | | | | | | | | | | | | **sub-30** | M | 35 | | | |
| **Mean** | **11F** | **23.62** | | **15F** | **66.57** | | **5F** | **33.91** | | **12F** | **27.21** | | **9F** | **21.35** | | **9F** | **56.13** | | **13F** | **28.42** |
| **SD** | | **2.96** | | | **7.39** | | | **11.32** | | | **3.73** | | | **2.98** | | | **17.01** | | | **10.01** |

**Table S2:** Gender and age characteristics of each participant involved in the *InLang* fMRI projects and database

#### MRI Acquisition

MRI data were collected using a whole-body 3 T MR scanner (Philips Achieva) with 40 mT/m gradient strength for all projects, except for Plast-LANG, which used a 3T MR scanner Bruker MedSpec S300 instead. For functional scans, the manufacturer-provided gradient-echo/T2* weighted EPI method was used.

##### NEREC

Forty-four adjacent axial slices parallel to the bicommissural plane were acquired in interleaved mode. Slice thickness was 3.5mm. The in-plane voxel size was 2.3 × 2.3 mm (220 × 220 × 131.75 mm field of view acquired with an 88 × 85 pixels data matrix; reconstructed with zero filling to 96 × 96 pixels). For the functional runs, the main sequence parameters were TR = 2.5 s, TE = 30 ms, and flip angle = 80°. A T1-weighted high-resolution (1 × 1 × 1mm) three-dimensional anatomical volume was also acquired.

##### NeuroMod

Forty-two adjacent axial slices parallel to the bicommissural plane were acquired in ascending mode. Slice thickness was 3 mm. During the functional run the cerebral volume was measured 192 times. The in-plane voxel size was 3 × 3 mm (240 × 240 mm field of view acquired with an 80 × 80 pixels data matrix; reconstructed with zero filling to 80 × 80 pixels). For the functional runs, the main sequence parameters were TR = 2.5 s, TE = 30 ms, and flip angle = 82°. A T1-weighted high-resolution (1 × 1 × 1mm) three-dimensional anatomical volume was also acquired.

##### Recovery

Fifty-two adjacent axial slices parallel to the bicommissural plane were acquired in non-interleaved ascendant mode. Acquisition parameters were: slice thickness 2.75 mm with a gap of 0.25 mm; in-plane voxel size 2.5 × 2.5 × 3 mm (220 × 220 × 156 mm field of view encoded with a 88 × 85 voxels matrix). For the functional runs, the main sequence parameters were TR = 3 sec, TE = 30 ms, flip angle = 80°. A T1-weighted high-resolution (1 × 1 × 1mm) three-dimensional anatomical volume was also acquired.

##### Plast-LANG

***Tasks: phonemic detection and semantic categorisation (PHON & SEM)***

Thirty-nine adjacent axial slices parallel to the bi-commissural plane were acquired in interleaved mode. Slice thickness was 3.5mm. During the functional run the cerebral volume was measured 155 times. The in-plane voxel size was 3 × 3 mm (216 × 216 mm field of view acquired with a 72 × 72 pixels data matrix; reconstructed with zero filling to 128 × 128 pixels). For the functional runs, the main sequence parameters were TR = 2.5 s, TE = 30 ms, and flip angle = 77°. A T1-weighted high-resolution (1 × 1 × 1mm) three-dimensional anatomical volume was also acquired.

***Task: prosodic judgment (PROS)***

Thirty-nine axial slices parallel to the anteroposterior commissural plane acquired in interleaved order (3 x 3 $mm^2$ in plane resolution with a slice thickness of 3.5 mm). During the functional run the cerebral volume was measured 173 times. For the functional runs, the main sequence parameters were TR = 3 s, TE = 40 ms, and flip angle = 77°. A T1-weighted high-resolution (1 × 1 × 1mm) three-dimensional anatomical volume was also acquired.

#### Reorg-GEREC

Forty-two adjacent axial slices parallel to the bicommissural plane were acquired in sequential mode. Slice thickness was 3 mm. The in-plane voxel size was 3 × 3 mm (field of view = 240 × 240 × 126 mm; data matrix = 80 × 80 pixels; reconstruction matrix = 80 × 80 pixels). For the functional runs, the main sequence parameters were TR = 2.5 s, TE = 30 ms, and flip angle = 82°. A T1-weighted high-resolution (1 × 1 × 1mm) three-dimensional anatomical volume was also acquired.

#### SEMVIE

Forty-four adjacent axial slices parallel to the bicommissural plane were acquired in interleaved mode. Slice thickness was 3.5mm. The in-plane voxel size was 2.3 × 2.3 mm (216 × 216 mm field of view acquired with a 72 × 72 pixels data matrix; reconstructed with zero filling to 128 × 128 pixels). For the functional runs, the main sequence parameters were TR = 2.5 s, TE = 30 ms, and flip angle = 77°. A T1-weighted high-resolution (1 × 1 × 1mm) three-dimensional anatomical volume was also acquired.

#### InnerSpeech

Forty-two adjacent axial slices parallel to the bi-commissural plane were acquired in non-interleaved mode. Slice thickness was 3 mm. The in-plane voxel size was 3 × 3 mm (240 × 240 mm field of view acquired with an 80 × 80 pixels data matrix). For the functional runs, the main sequence parameters were TR = 2.5 s, TE = 30 ms, and flip angle = 82°. A T1-weighted high-resolution (1 × 1 × 1mm) three-dimensional anatomical volume was also acquired.

#### Data Analyses

##### Functional MRI Processing

###### *Pre-processing*

Data analysis was performed by using the general linear model (Friston et al., 1995) in SPM12 (Wellcome Department of Imaging Neuroscience, London, UK, www.fil.ion.ucl.ac.uk/spm) implemented in MATLAB (Mathworks Inc., Sherborn, MA, USA). First, functional volumes were time-corrected with regard to the following slice as a reference: 22$^{nd}$ (NEREC and SEMVIE), 21$^{st}$ (NeuroMod, Reorg-GEREC and InnerSpeech), 26$^{th}$ (Recovery), 19$^{th}$ (Plast-LANG). All volumes were then realigned to correct for head motion using rigid body transformations (3 translations and 3 rotations). The T1-weighted anatomical volume was coregistered to mean images created by the realignment procedure and was normalised to the MNI space using a trilinear interpolation. The spatial resolution of the anatomical volume was of 1 x 1 x 1 mm. The anatomical normalisation parameters were subsequently used for the normalisation of functional volumes. Finally, each functional volume was smoothed by a 6 mm FWHM (Full Width at Half Maximum) Gaussian kernel. Time series for each voxel were high-pass filtered to remove low-frequency noise and signal drift with the following cut-offs: 1/384 Hz (NEREC), 1/256 Hz (NeuroMod), 1/128 Hz (Recovery, Plast-LANG, Reorg-GEREC and SEMVIE) and 1/512 Hz (InnerSpeech).

###### *Statistical analyses: individual activation maps (1$^{st}$ level)*

Statistical analyses were performed separately by project and task. For each task, regressors have been created to match conditions of interest. The object naming tasks (regardless of project) and all tasks in the InnerSpeech project were analysed with a "task" regressor vs implicit baseline. For all other cases, contrasts involving "task" vs "control" were set up. We note that although a control condition was available in some of the projects for the object naming task, we analysed all cases with a "task" regressor vs implicit baseline for uniformity across projects. These regressors were convolved with a canonical hemodynamic response function (HRF). The movement parameters derived from the realignment corrections (3 translations and 3 rotations) were included in the design matrix as nuisance regressors. The general linear model was then used to generate the parameter estimates of activity for each voxel, each condition and each participant. The statistical parametric maps were generated from linear contrasts between the HRF parameter estimates for the conditions of interest. The spatial resolution of statistical parametric maps was of 3 x 3 x 3 mm. The statistical analyses have been performed at the individual level to calculate contrasts of task *vs* control or task *vs* baseline, as appropriate, in order to highlight brain regions specifically involved with the following main linguistic processes: semantic processing, phonology/prosody, lexical or lexico-syntactic production, dialogue and social aspects of language, monitoring or even spontaneous speech (see Table S1 above).

##### *Statistical analyses: 2nd level and clustering based on the similarity of BOLD activations*

We have performed the second level statistical analyses for all tasks separately. The resulting statistical fMRI maps (FDR corr., k =5) were mapped onto the surface using FreeSurfer (fsaverage). We then extracted the t values on the vertex-wise Conte-69 standard cortical surface mesh (32k vertices; Van Essen et al., 2012). A vector of 32.000 indexed t-values containing the spatial location information was obtained for each task. We have binarized the vectors (0 for vertices below the statistical threshold and 1 otherwise). We then computed a task-based similarity matrix from the Jaccard indices calculated between the respective vectors (similarity = 1 - the Jaccard distance; Figure S1).

Finally, we applied data-driven hierarchical clustering. We observed clustering into 5 main groups, based on the similarity of neurofunctional BOLD activations. The nature of the grouped tasks, with respect to the linguistic subprocesses primarily targeted by the protocols (Table S1), allowed us to label the task-groups identified based on data. Accordingly, we propose the following designation: DECODING, SEMANTIC, PRODUCTION, MONITORING, and WANDERING (see also Figure 2A in the main text).

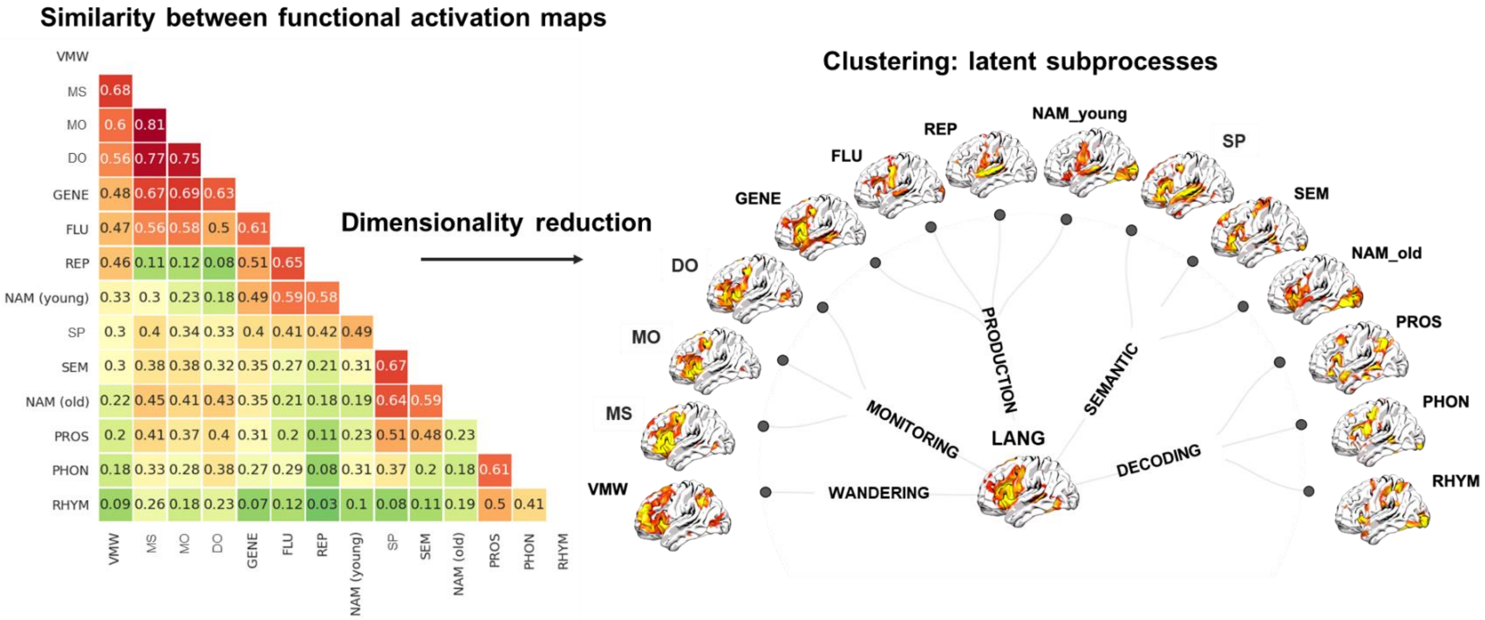


**Figure S1:** Similarity of thresholded activation maps and assumed underlying subprocesses.
