## Appendix S2 for "Unraveling the functional attributes of the language connectome: crucial subnetworks, flexibility and variability"

### Supplementary information (SI) – Results

### LANG: task-related connectomic atlas

#### Global properties

**Table S3**: Global metrics collected on the networks of each task and subprocesses.

Nb_nodes: number of non-isolated nodes in the subnetwork composed of top 5% of positive edges; Mean_Rval: average correlation values for the entire (thresholded) network; Ndensity: density of network nodes relative to the average number of nodes selected from a distribution of random networks (n = 100) and composed of the same number of edges (fixed number of 3485 edges; 5%); Eglob: global efficiency measured at the network level; Eloc: local efficiency calculated at the network level; I:S balance: relative difference between Eglob (Integration) and Eloc (Segregation); Mean_Geodist_mm: average measure of the existing geodesic distance between interconnected nodes of the network (estimated in millimeters); Cluster_Geo_dist: clustering according to the global connectivity profile accounting for geodesic distance, which could be interpreted as: 1 = short-range; 2 = middle-range; 3 = long-range functional connectivity. See the Materials and Methods Section in the main text for technical details of global metrics (Section 5.3.1.). See Appendix S1 for the description of tasks and subprocesses.

| Variables | Nb_nodes | Mean_Rval | Ndensity | Eglob | Eloc | I:S balance | Mean_ Geodist_mm | Cluster_ Geo_dist |
|---|---|---|---|---|---|---|---|---|
| **MONITORING** | 109 | 0.541 | 0.757 | 0.622 | 0.832 | -0.21 | 15.7 | 1 |
| MS | 112 | 0.524 | 0.778 | 0.586 | 0.804 | -0.217 | 18.1 | 1 |
| MO | 110 | 0.542 | 0.764 | 0.624 | 0.834 | -0.209 | 25.2 | 1 |
| DO | 108 | 0.536 | 0.75 | 0.641 | 0.837 | -0.196 | 28.9 | 1 |
| **DECODING** | 103 | 0.596 | 0.715 | 0.501 | 0.846 | -0.345 | 31.7 | 1 |
| PHON | 100 | 0.621 | 0.694 | 0.533 | 0.876 | -0.343 | 36.7 | 2 |
| PROS | 105 | 0.568 | 0.729 | 0.49 | 0.863 | -0.373 | 27.9 | 1 |
| RHYM | 104 | 0.587 | 0.722 | 0.522 | 0.827 | -0.305 | 37.6 | 2 |
| **SEMANTIC** | 113 | 0.617 | 0.785 | 0.66 | 0.619 | 0.041 | 63.7 | 3 |
| SEM | 113 | 0.618 | 0.785 | 0.684 | 0.59 | 0.094 | 68.9 | 3 |
| SP | 109 | 0.621 | 0.757 | 0.659 | 0.614 | 0.045 | 62.6 | 3 |
| **PRODUCTION** | 120 | 0.492 | 0.833 | 0.619 | 0.581 | 0.038 | 42.3 | 2 |
| NAM | 121 | 0.513 | 0.84 | 0.608 | 0.605 | 0.004 | 52.2 | 2 |
| FLU | 127 | 0.497 | 0.882 | 0.595 | 0.562 | 0.033 | 43.5 | 2 |
| GENE | 119 | 0.456 | 0.826 | 0.637 | 0.618 | 0.018 | 51.7 | 2 |
| REP | 114 | 0.519 | 0.792 | 0.635 | 0.62 | 0.015 | 39.7 | 2 |
| **WANDERING** | 125 | 0.443 | 0.868 | 0.82 | 0.704 | 0.116 | 76.8 | 3 |
| VMW | 125 | 0.443 | 0.868 | 0.82 | 0.704 | 0.116 | 76.8 | 3 |
| **LANG** | 131 | 0.481 | 0.930 | 0.653 | 0.604 | 0.049 | 68.1 | 3 |

#### The LANG atlas, Nets and nodal properties

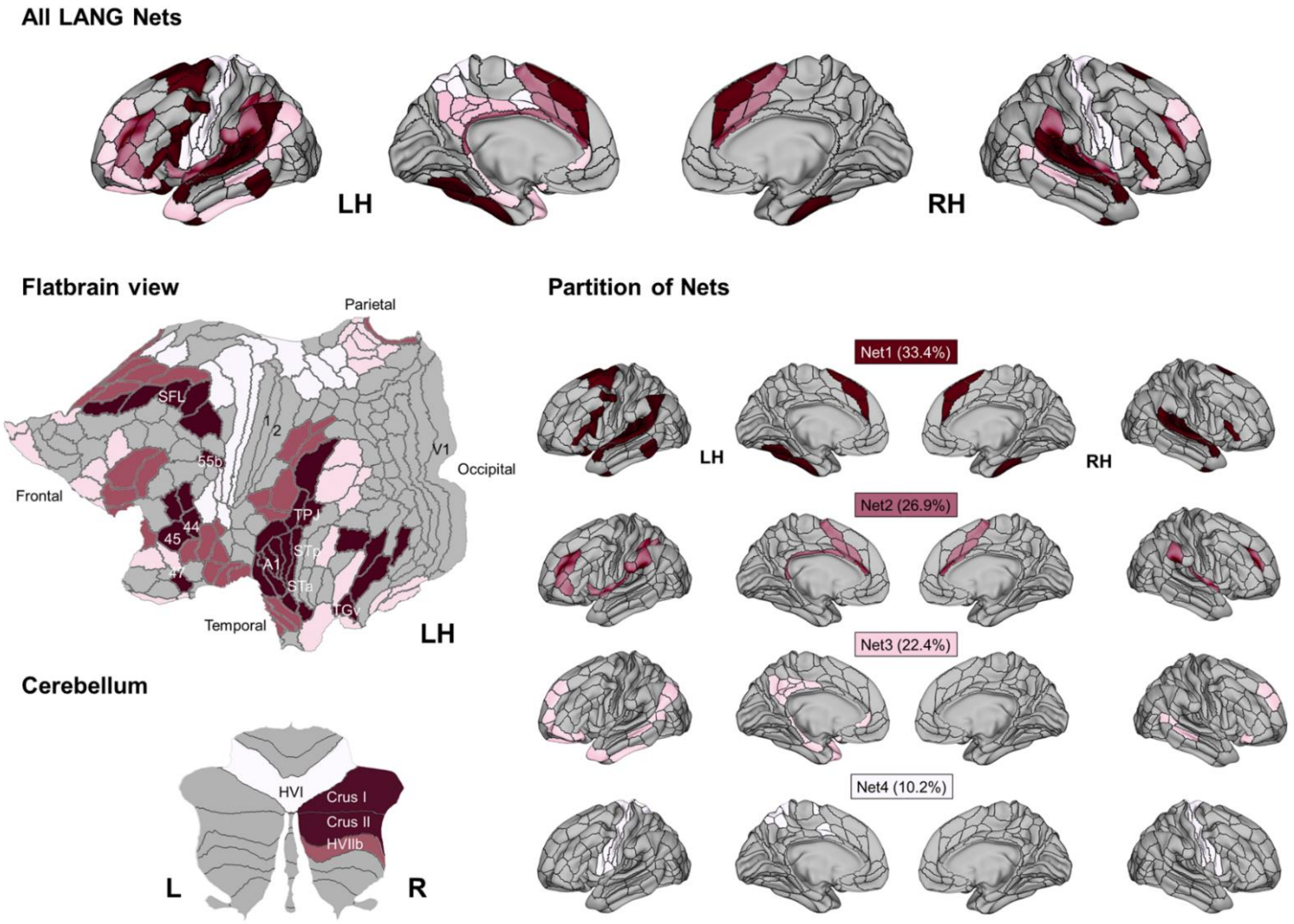

**Figure S2:** Illustration of the LANG connectomic atlas projected on the two hemispheres, flatbrain and cerebellum.

The LANG Nets distribution corresponds to the Glasser HCP parcellation (Glasser et al., 2016). The distribution on the basis of Power's brain coordinates (Power et al., 2011) is slightly different (Net 1 = 41.2%; Net 2 = 21.4%; Net 3 = 19.8%; Net 4 = 17.6%; Table S4). LH = left hemisphere; RH = right hemisphere.

**The LANG atlas package** (containing the .nodes files of All ROI peaks, LH peaks, RH peaks, cerebellum peaks, and the LANG matrices) is downloadable here:

```
https://doi.org/10.5281/zenodo.6402396
```

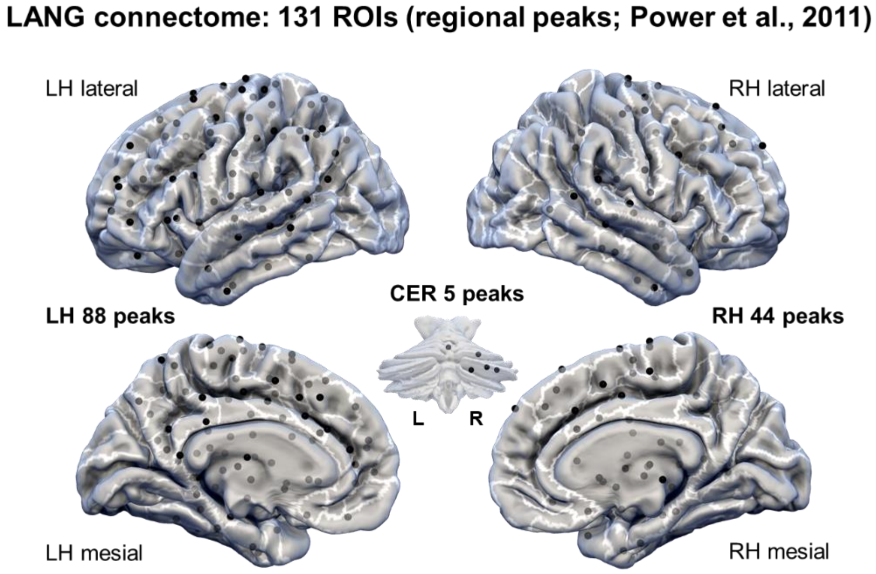

**Table S4**: Functional information about LANG ROIs (local connectivity)

Brain x, y, z coordinates (Power et al., 2011) of all the LANG regions of interest (n = 131 ROIs). The LANG ROIs are sorted by Nets.

Name_label: labeling of the ROIs brain peaks using the AICHA atlas (Joliot et al., 2015), allowing a fine and homotopic functional labeling; Glasser_assign: labeling of LANG ROIs based on Glasser's proposed multimodal parcellation (version 1.0: HCP_MMP1.0 proposed by Glasser et al. 2016); CAB_NP_assign: labeling of LANG ROIs according to the Cole-Anticevic brain-wide functional partition (CAB-NP; Ji et al., 2019); Lang_Net_assign: assignment to a specific community, following the modularity analyses applied on the LANG general connectome deriving from the set of tasks (*InLang* database). 1 = Net1, 2 = Net2, 3 = Net3, 4 = Net4; Clustering_coeff: clustering coefficient; DC_i: degree centrality of a ROI in the LANG general connectome; DC_ki: degree centrality or "k" parameter corresponding to the degree of interconnection of a given ROI with the other ROIs of its LANG Net; zi: within-component degree z-score (intra-Net FC); PCi: participation coefficient (inter-Net FC); zPCi: participation coefficient z-scored; Func_role: zi and zPCi have been used to assign a functional role of each of the LANG ROI: connector (high $z_i$/high $zPC_i$; high intra-Net and high inter-Net FC); provincial (high $z_i$/low $zPC_i$; high intra-Net FC); satellite (high $z_i$/low $zPC_i$; high inter-Net FC); or peripheral (low *zi*/low *zPCi*; low inter-Net FC). See the Materials and Methods Section in the main text for technical aspects about local metrics (Section 5.3.4.).

| Coord_Power | | | | | | | | | | | | | |
|---|---|---|---|---|---|---|---|---|---|---|---|---|---|
| x | y | z | Name_label (AICHA atlas) | Glasser_assign | CAB_NP_ assign | LANG_ Net_assign | Clustering _coeff | DCi | DC_ ki | zi | PCi | z_Pci | Func_role |
| 22 | -58 | -23 | **Cerebelum_6_R** | NaN | NaN | 1 | 0.748 | 0.469 | 16 | -1.023 | 0.931 | 0.73 | Satellite |
| 28 | -77 | -32 | **Cerebelum_Crus1_R_1** | NaN | NaN | 1 | 0.682 | 0.408 | 24 | 0.842 | 0.795 | 0.144 | Connector |
| 35 | -67 | -34 | **Cerebelum_Crus1_R_2** | NaN | NaN | 1 | 0.641 | 0.246 | 25 | 1.075 | 0.39 | -1.598 | Provincial |
| 0 | 30 | 27 | **Cingulum_Ant_L_5** | d32/p24 | Default | 1 | 0.713 | 0.592 | 17 | -0.79 | 0.951 | 0.816 | Satellite |
| 5 | 23 | 37 | **Cingulum_Mid_R_3** | d32 | CON | 1 | 0.695 | 0.369 | 19 | -0.324 | 0.843 | 0.352 | Satellite |
| -54 | 22 | 21 | **Frontal_Inf_Tri_L_2** | 45 | Language | 1 | 0.641 | 0.677 | 30 | 2.241 | 0.884 | 0.526 | Connector |
| -49 | 25 | -1 | **Frontal_Inf_Tri_L_3** | 44 | Language | 1 | 0.908 | 0.423 | 20 | -0.091 | 0.868 | 0.457 | Satellite |
| 48 | 22 | 10 | **Frontal_Inf_Tri_R_2** | 45 | Language | 1 | 0.481 | 0.308 | 20 | -0.091 | 0.75 | -0.049 | Peripheral |
| -42 | 38 | 21 | **Frontal_Mid_L_2** | IFSp/IFja | Language | 1 | 0.581 | 0.208 | 24 | 0.842 | 0.21 | -2.371 | Provincial |
| -16 | -5 | 71 | **Frontal_Sup_L_1** | SFL | Language | 1 | 0.626 | 0.608 | 22 | 0.376 | 0.922 | 0.692 | Connector |
| -23 | 11 | 64 | **Frontal_Sup_L_6** | 6a | DAN | 1 | 0.664 | 0.646 | 22 | 0.376 | 0.931 | 0.73 | Connector |
| -3 | 26 | 44 | **Frontal_Sup_Medial_L_4** | 8BM | FPN | 1 | 0.629 | 0.638 | 22 | 0.376 | 0.93 | 0.723 | Connector |
| 13 | 30 | 59 | **Frontal_Sup_Medial_R_4** | 8BM | FPN | 1 | 0.807 | 0.431 | 24 | 0.842 | 0.816 | 0.236 | Connector |
| 19 | -8 | 64 | **Frontal_Sup_R_1** | SFL | Language | 1 | 0.728 | 0.246 | 15 | -1.257 | 0.78 | 0.081 | Satellite |
| -37 | -29 | -26 | **Fusiform_L_1** | VVC | Default | 1 | 0.649 | 0.5 | 18 | -0.557 | 0.923 | 0.696 | Satellite |
| -34 | -38 | -16 | **Fusiform_L_2** | PHA3 | Default | 1 | 0.714 | 0.562 | 16 | -1.023 | 0.952 | 0.819 | Satellite |
| -31 | -10 | -36 | **Fusiform_L_3** | TF/TE2a | Default | 1 | 0.736 | 0.562 | 24 | 0.842 | 0.892 | 0.561 | Connector |
| 33 | -12 | -34 | **Fusiform_R_3** | TF | Default | 1 | 0.71 | 0.162 | 14 | -1.49 | 0.556 | -0.885 | Peripheral |
| -31 | 19 | -19 | **Insula_L_1** | 47s | Default | 1 | 0.555 | 0.585 | 16 | -1.023 | 0.956 | 0.835 | Satellite |
| -53 | -49 | 43 | **Parietal_Inf_L_2** | PFm | FPN | 1 | 0.477 | 0.723 | 29 | 2.008 | 0.905 | 0.616 | Connector |
| -32 | -1 | 54 | **Precentral_L_6** | 55b | Language | 1 | 0.574 | 0.723 | 29 | 2.008 | 0.905 | 0.616 | Connector |
| -22 | 7 | -5 | **Putamen_L_1** | NaN | NaN | 1 | 0.629 | 0.623 | 23 | 0.609 | 0.919 | 0.679 | Connector |
| 31 | -14 | 2 | **Putamen_R_1** | NaN | NaN | 1 | 0.554 | 0.308 | 17 | -0.79 | 0.819 | 0.249 | Satellite |
| -45 | 0 | 9 | **Rolandic_Oper_L_1** | Ig | Auditory | 1 | 0.604 | 0.508 | 18 | -0.557 | 0.926 | 0.706 | Satellite |
| -38 | -33 | 17 | **Rolandic_Oper_L_2** | RI | Auditory | 1 | 0.781 | 0.546 | 24 | 0.842 | 0.886 | 0.534 | Connector |
| -55 | -9 | 12 | **Rolandic_Oper_L_3** | OP2-3 | Auditory | 1 | 0.891 | 0.085 | 10 | -2.422 | 0.174 | -2.527 | Peripheral |
| 49 | 8 | -1 | **Rolandic_Oper_R_1** | Ig | Auditory | 1 | 0.635 | 0.215 | 18 | -0.557 | 0.587 | -0.751 | Peripheral |
| 43 | -23 | 20 | **Rolandic_Oper_R_2** | RI | Auditory | 1 | 0.752 | 0.277 | 23 | 0.609 | 0.592 | -0.729 | Provincial |
| 56 | -5 | 13 | **Rolandic_Oper_R_3** | OP2-3 | Auditory | 1 | 0.601 | 0.215 | 16 | -1.023 | 0.673 | -0.378 | Peripheral |
| -10 | 11 | 67 | **Supp_Motor_Area_L_3** | SFL | Language | 1 | 0.644 | 0.354 | 21 | 0.142 | 0.792 | 0.129 | Connector |
| -1 | 15 | 44 | **Supp_Motor_Area_L_4** | 8BM | FPN | 1 | 0.571 | 0.115 | 12 | -1.956 | 0.36 | -1.726 | Peripheral |
| 3 | -17 | 58 | **Supp_Motor_Area_R_3** | SFL | Language | 1 | 0.692 | 0.546 | 24 | 0.842 | 0.886 | 0.534 | Connector |
| 13 | -1 | 70 | **Supp_Motor_Area_R_4** | 8BM | FPN | 1 | 0.536 | 0.338 | 17 | -0.79 | 0.851 | 0.384 | Satellite |
| -60 | -25 | 14 | **SupraMarginal_L_1** | STV | Language | 1 | 0.57 | 0.708 | 28 | 1.775 | 0.907 | 0.627 | Connector |
| -50 | -34 | 26 | **SupraMarginal_L_2** | PSL | Language | 1 | 0.511 | 0.554 | 20 | -0.091 | 0.923 | 0.694 | Satellite |
| -50 | -7 | -39 | **Temporal_Inf_L_2** | TGv | Language | 1 | 0.745 | 0.562 | 21 | 0.142 | 0.917 | 0.67 | Connector |
| -47 | -51 | -21 | **Temporal_Inf_L_3** | TE2p | Default | 1 | 0.598 | 0.669 | 15 | -1.257 | 0.97 | 0.898 | Satellite |
| 49 | -3 | -38 | **Temporal_Inf_R_5** | TGv | Language | 1 | 0.441 | 0.292 | 20 | -0.091 | 0.723 | -0.165 | Peripheral |
| -56 | -13 | -10 | **Temporal_Mid_L_4** | STSda | Language | 1 | 0.691 | 0.131 | 16 | -1.023 | 0.114 | -2.782 | Peripheral |
| -58 | -30 | -4 | **Temporal_Mid_L_5** | TPOJ1 | Language | 1 | 0.707 | 0.585 | 23 | 0.609 | 0.908 | 0.632 | Connector |
| -68 | -41 | -5 | **Temporal_Mid_L_6** | TE1P | FPN | 1 | 0.741 | 0.515 | 18 | -0.557 | 0.928 | 0.715 | Satellite |
| 52 | -2 | -16 | **Temporal_Mid_R_4** | STSda | Language | 1 | 0.553 | 0.208 | 22 | 0.376 | 0.336 | -1.829 | Provincial |
| 51 | -29 | -4 | **Temporal_Mid_R_5** | TPOJ1 | Language | 1 | 0.604 | 0.285 | 16 | -1.023 | 0.813 | 0.221 | Satellite |
| 46 | 16 | -30 | **Temporal_Pole_Mid_R_1** | STGa | Language | 1 | 0.51 | 0.192 | 17 | -0.79 | 0.538 | -0.962 | Peripheral |
| -51 | 8 | -2 | **Temporal_Pole_Sup_L** | STGa | Language | 1 | 0.698 | 0.423 | 25 | 1.075 | 0.793 | 0.137 | Connector |
| -49 | -26 | 5 | **Temporal_Sup_L_1** | MBelt/A1 | Auditory | 1 | 0.587 | 0.454 | 24 | 0.842 | 0.835 | 0.314 | Connector |
| -55 | -40 | 14 | **Temporal_Sup_L_2** | PBelt/A4/A5 | Auditory | 1 | 0.595 | 0.708 | 21 | 0.142 | 0.948 | 0.801 | Connector |
| 65 | -33 | 20 | **Temporal_Sup_R_1** | PBelt/A4 | Auditory | 1 | 0.717 | 0.408 | 21 | 0.142 | 0.843 | 0.35 | Connector |
| 58 | -16 | 7 | **Temporal_Sup_R_2** | MBelt/R52 | Auditory | 1 | 0.708 | 0.231 | 16 | -1.023 | 0.716 | -0.197 | Peripheral |
| 52 | -33 | 8 | **Temporal_Sup_R_4** | A5 | Language | 1 | 0.696 | 0.3 | 20 | -0.091 | 0.737 | -0.105 | Peripheral |
| 56 | -46 | 11 | **Temporal_Sup_R_5** | STV/PSL | Language | 1 | 0.783 | 0.477 | 22 | 0.376 | 0.874 | 0.484 | Connector |
| -2 | -13 | 12 | **Thalamus_L_1** | NaN | NaN | 1 | 0.652 | 0.2 | 25 | 1.075 | 0.075 | -2.949 | Provincial |
| 12 | -17 | 8 | **Thalamus_R_2** | NaN | NaN | 1 | 0.447 | 0.477 | 20 | -0.091 | 0.896 | 0.578 | Satellite |
| 9 | -4 | 6 | **Thalamus_Ven_Ant_Nucleus_R** | NaN | NaN | 1 | 0.508 | 0.354 | 22 | 0.376 | 0.771 | 0.042 | Connector |

**Coord_Power**

| x | y | z | Name_label (AICHA atlas) | Glasser_assign | CAB_NP_ assign | LANG_ Net_assign | Clustering _coeff | DCi | DC_ ki | zi | PCi | z_Pci | Func_role |
|---|---|---|---|---|---|---|---|---|---|---|---|---|---|
| 17 | -80 | -34 | **Cerebelum_Crus2_R** | NaN | NaN | 2 | 0.629 | 0.469 | 16 | -1.704 | 0.931 | 0.578 | Satellite |
| -3 | 42 | 16 | **Cingulum_Ant_L_3** | p24/a24pr | CON | 2 | 0.695 | 0.208 | 13 | -1.858 | 0.768 | -1.284 | Peripheral |
| -11 | 26 | 25 | **Cingulum_Ant_L_4** | 33pr | CON | 2 | 0.848 | 0.469 | 18 | -1.602 | 0.913 | 0.369 | Satellite |
| 10 | 22 | 27 | **Cingulum_Ant_R_2** | p24/a24pr | CON | 2 | 0.547 | 0.254 | 18 | -1.602 | 0.702 | -2.034 | Peripheral |
| -5 | 18 | 34 | **Cingulum_Mid_L_2** | a32pr | Default | 2 | 0.718 | 0.585 | 14 | -1.807 | 0.966 | 0.976 | Satellite |
| -2 | -35 | 31 | **Cingulum_Post_L** | RSC | FPN | 2 | 0.652 | 0.269 | 18 | -1.602 | 0.736 | -1.657 | Peripheral |
| -35 | 20 | 51 | **Frontal_Mid_L_1** | 9-46d | CON | 2 | 0.847 | 0.446 | 9 | -2.063 | 0.976 | 1.089 | Satellite |
| -34 | 55 | 4 | **Frontal_Mid_L_3** | p47r | FPN | 2 | 0.651 | 0.638 | 9 | -2.063 | 0.988 | 1.229 | Satellite |
| -28 | 52 | 21 | **Frontal_Mid_L_4** | 946v/46 | FPN | 2 | 0.783 | 0.508 | 24 | -1.294 | 0.868 | -0.146 | Peripheral |
| -39 | 51 | 17 | **Frontal_Mid_L_5** | a946v | FPN | 2 | 0.69 | 0.554 | 14 | -1.807 | 0.962 | 0.932 | Satellite |
| 22 | 39 | 39 | **Frontal_Sup_R_3** | 9-46d | CON | 2 | 0.733 | 0.115 | 8 | -2.114 | 0.716 | -1.885 | Peripheral |
| -35 | 20 | 0 | **Insula_L_2** | MI/Pol2 | CON | 2 | 0.494 | 0.315 | 10 | -2.012 | 0.941 | 0.684 | Satellite |
| 27 | 16 | -17 | **Insula_R_2** | MI/Pol2 | CON | 2 | 0.562 | 0.377 | 19 | -1.55 | 0.85 | -0.353 | Peripheral |
| 37 | 1 | -4 | **Insula_R_6** | Pol1/Pol2 | CON | 2 | 0.671 | 0.162 | 10 | -2.012 | 0.773 | -1.226 | Peripheral |
| -54 | -23 | 43 | **Parietal_Inf_L_1** | PF/Pfop | CON | 2 | 0.592 | 0.4 | 15 | -1.755 | 0.917 | 0.413 | Satellite |
| -28 | -58 | 48 | **Parietal_Inf_L_3** | IP1 | FPN | 2 | 0.521 | 0.292 | 16 | -1.704 | 0.823 | -0.661 | Peripheral |
| -33 | -46 | 47 | **Parietal_Inf_L_5** | IP2 | FPN | 2 | 0.636 | 0.262 | 11 | -1.96 | 0.895 | 0.168 | Satellite |
| 49 | -42 | 45 | **Parietal_Inf_R_1** | PF/Pfop | CON | 2 | 0.785 | 0.492 | 12 | -1.909 | 0.965 | 0.962 | Satellite |
| -45 | -32 | 47 | **Postcentral_L_3** | AIP | DAN | 2 | 0.612 | 0.377 | 13 | -1.858 | 0.93 | 0.56 | Satellite |
| -13 | -40 | 1 | **Precuneus_L_2** | RSC | FPN | 2 | 0.643 | 0.246 | 14 | -1.807 | 0.809 | -0.822 | Peripheral |
| -3 | 2 | 53 | **Supp_Motor_Area_L_2** | a32pr/p32pr/SCEF | CON | 2 | 0.578 | 0.238 | 13 | -1.858 | 0.824 | -0.645 | Peripheral |
| 10 | -2 | 45 | **Supp_Motor_Area_R_1** | a32pr/p32pr/SCEF | CON | 2 | 0.62 | 0.146 | 9 | -2.063 | 0.776 | -1.199 | Peripheral |
| 10 | -17 | 74 | **Supp_Motor_Area_R_2** | SCEF | CON | 2 | 0.758 | 0.492 | 20 | -1.499 | 0.902 | 0.248 | Satellite |
| 7 | 8 | 51 | **Supp_Motor_Area_R_5** | p32pr | CON | 2 | 0.73 | 0.492 | 17 | -1.653 | 0.929 | 0.558 | Satellite |
| -53 | -22 | 23 | **SupraMarginal_L_3** | PFcm | CON | 2 | 0.81 | 0.485 | 9 | -2.063 | 0.98 | 1.131 | Satellite |
| 55 | -45 | 37 | **SupraMarginal_R_2** | PFcm | CON | 2 | 0.859 | 0.477 | 16 | -1.704 | 0.933 | 0.603 | Satellite |
| -10 | -18 | 7 | **Thalamus_L_2** | NaN | NaN | 2 | 0.698 | 0.615 | 11 | -1.96 | 0.981 | 1.148 | Satellite |
| 6 | -24 | 0 | **Thalamus_R_1** | NaN | NaN | 2 | 0.685 | 0.223 | 9 | -2.063 | 0.904 | 0.264 | Satellite |

**Coord_Power**

| x | y | z | Name_label (AICHA atlas) | Glasser_assign | CAB_NP_ assign | LANG_ Net_assign | Clustering _coeff | DCi | DC_ ki | zi | PCi | z_Pci | Func_role |
|---|---|---|---|---|---|---|---|---|---|---|---|---|---|
| -44 | -65 | 35 | **Angular_L_1** | PGi | Default | 3 | 0.517 | 0.3 | 12 | -2.574 | 0.905 | 0.041 | Satellite |
| -11 | 45 | 8 | **Cingulum_Ant_L_2** | a24 | Default | 3 | 0.685 | 0.585 | 15 | -2.408 | 0.961 | 0.336 | Satellite |
| -10 | -2 | 42 | **Cingulum_Mid_L_1** | 23d | Default | 3 | 0.529 | 0.162 | 9 | -2.741 | 0.816 | -0.43 | Peripheral |
| -2 | -37 | 44 | **Cingulum_Mid_L_3** | 31a/31pv/31pd | Default | 3 | 0.667 | 0.031 | 4 | -3.02 | 0,000 | -4.745 | Peripheral |
| 49 | 35 | -12 | **Frontal_Inf_Orb_R_2** | 47l/OFC | Default | 3 | 0.585 | 0.315 | 18 | -2.241 | 0.807 | -0.478 | Peripheral |
| -47 | 11 | 23 | **Frontal_Inf_Tri_L_1** | 47l | Default | 3 | 0.699 | 0.515 | 13 | -2.519 | 0.962 | 0.342 | Satellite |
| -21 | 41 | -20 | **Frontal_Mid_Orb_L_1** | 47l/OFC | Default | 3 | 0.582 | 0.508 | 15 | -2.408 | 0.948 | 0.268 | Satellite |
| -42 | 45 | -2 | **Frontal_Mid_Orb_L_2** | a47r | Default | 3 | 0.825 | 0.508 | 13 | -2.519 | 0.961 | 0.336 | Satellite |
| -16 | 29 | 53 | **Frontal_Sup_L_2** | 9p | Default | 3 | 0.683 | 0.469 | 16 | -2.352 | 0.931 | 0.178 | Satellite |
| -20 | 45 | 39 | **Frontal_Sup_L_4** | 9a | Default | 3 | 0.653 | 0.646 | 10 | -2.686 | 0.986 | 0.467 | Satellite |
| 23 | 33 | 48 | **Frontal_Sup_R_2** | 9p | Default | 3 | 0.765 | 0.492 | 19 | -2.185 | 0.912 | 0.075 | Satellite |
| 13 | 55 | 38 | **Frontal_Sup_R_4** | 9a | Default | 3 | 0.826 | 0.485 | 12 | -2.574 | 0.964 | 0.35 | Satellite |
| -21 | -22 | -20 | **ParaHippocampal_L_1** | PreS | Default | 3 | 0.681 | 0.577 | 10 | -2.686 | 0.982 | 0.447 | Satellite |
| -26 | -40 | -8 | **ParaHippocampal_L_2** | EC | Default | 3 | 0.562 | 0.362 | 14 | -2.463 | 0.911 | 0.072 | Satellite |
| -42 | -55 | 45 | **Parietal_Inf_L_4** | PGs | Default | 3 | 0.507 | 0.523 | 18 | -2.241 | 0.93 | 0.171 | Satellite |
| -7 | -55 | 27 | **Precuneus_L_3** | 31pd | Default | 3 | 0.697 | 0.615 | 11 | -2.63 | 0.981 | 0.442 | Satellite |
| -11 | -56 | 16 | **Precuneus_L_4** | v23ab | Default | 3 | 0.866 | 0.446 | 13 | -2.519 | 0.95 | 0.276 | Satellite |
| -3 | -49 | 13 | **Precuneus_L_5** | 31pv/d23ab | Default | 3 | 0.881 | 0.431 | 18 | -2.241 | 0.897 | -0.005 | Peripheral |
| -7 | -71 | 42 | **Precuneus_L_6** | 7m | Default | 3 | 0.716 | 0.562 | 13 | -2.519 | 0.968 | 0.374 | Satellite |
| 55 | -31 | -17 | **Temporal_Inf_R_2** | STSvp | Default | 3 | 0.616 | 0.354 | 18 | -2.241 | 0.847 | -0.268 | Peripheral |
| -46 | -61 | 21 | **Temporal_Mid_L_1** | TPOJ3 | Multi | 3 | 0.814 | 0.477 | 16 | -2.352 | 0.933 | 0.189 | Satellite |
| -53 | 3 | -27 | **Temporal_Mid_L_7** | TGd | Default | 3 | 0.658 | 0.415 | 10 | -2.686 | 0.966 | 0.36 | Satellite |
| -49 | -42 | 1 | **Temporal_Mid_L_8** | STSvp | Default | 3 | 0.719 | 0.554 | 14 | -2.463 | 0.962 | 0.342 | Satellite |
| -56 | -50 | 10 | **Temporal_Mid_L_9** | TPOJ2 | Multi | 3 | 0.76 | 0.338 | 12 | -2.574 | 0.926 | 0.148 | Satellite |
| 65 | -24 | -19 | **Temporal_Mid_R_1** | STSvp | Default | 3 | 0.784 | 0.492 | 12 | -2.574 | 0.965 | 0.356 | Satellite |
| 46 | -59 | 4 | **Temporal_Mid_R_6** | TPOJ2 | Multi | 3 | 0.833 | 0.492 | 12 | -2.574 | 0.965 | 0.356 | Satellite |

| Coord_Power | | | | | | | | | | | | | |
|---|---|---|---|---|---|---|---|---|---|---|---|---|---|
| x | y | z | Name_label (AICHA atlas) | Glasser_assign | CAB_NP_ assign | LANG_ Net_assign | Clustering_ coeff | DCi | DC_ ki | zi | PCi | z_Pci | Func_role |
| -7 | -21 | 65 | **Paracentral_Lobule_L_1** | 6mp | SM | 4 | 0.562 | 0.292 | 6 | -1.535 | 0.975 | 0.871 | Satellite |
| -7 | -33 | 72 | **Paracentral_Lobule_L_2** | 5L | SM | 4 | 0.691 | 0.131 | 16 | 0.867 | 0.114 | -1.734 | Provincial |
| -13 | -17 | 75 | **Paracentral_Lobule_L_3** | 6mp | SM | 4 | 0.762 | 0.169 | 9 | -0.814 | 0.833 | 0.44 | Satellite |
| -21 | -31 | 61 | **Postcentral_L_4** | 3b | SM | 4 | 0.733 | 0.346 | 12 | -0.094 | 0.929 | 0.732 | Satellite |
| -49 | -11 | 35 | **Postcentral_L_5** | 3a | SM | 4 | 0.56 | 0.108 | 11 | -0.334 | 0.383 | -0.921 | Peripheral |
| -53 | -10 | 24 | **Postcentral_L_6** | 1 | SM | 4 | 0.733 | 0.115 | 15 | 0.626 | 0,000 | -2.08 | Provincial |
| 51 | -6 | 32 | **Postcentral_R_5** | 3a | SM | 4 | 0.701 | 0.208 | 19 | 1.587 | 0.505 | -0.552 | Provincial |
| 66 | -8 | 25 | **Postcentral_R_6** | 1 | SM | 4 | 0.477 | 0.4 | 10 | -0.574 | 0.963 | 0.835 | Satellite |
| -38 | -15 | 69 | **Precentral_L_1** | 4 | SM | 4 | 0.576 | 0.469 | 8 | -1.055 | 0.983 | 0.895 | Satellite |
| -40 | -19 | 54 | **Precentral_L_2** | 4 | SM | 4 | 0.733 | 0.115 | 10 | -0.574 | 0.556 | -0.398 | Peripheral |
| -38 | -27 | 69 | **Precentral_L_3** | 4 | SM | 4 | 0.56 | 0.108 | 10 | -0.574 | 0.49 | -0.597 | Peripheral |
| -44 | 2 | 46 | **Precentral_L_4** | 3a | SM | 4 | 0.565 | 0.5 | 11 | -0.334 | 0.931 | 0.739 | Satellite |
| -41 | 6 | 33 | **Precentral_L_5** | 6v | SM | 4 | 0.537 | 0.277 | 10 | -0.574 | 0.981 | 0.889 | Satellite |
| 29 | -17 | 71 | **Precentral_R_1** | 4 | SM | 4 | 0.456 | 0.238 | 19 | 1.587 | 0.624 | -0.19 | Provincial |
| 44 | -8 | 57 | **Precentral_R_2** | 4 | SM | 4 | 0.733 | 0.115 | 12 | -0.094 | 0.36 | -0.99 | Peripheral |
| 42 | -20 | 55 | **Precentral_R_3** | 4 | SM | 4 | 0.685 | 0.223 | 18 | 1.347 | 0.615 | -0.219 | Provincial |
| 47 | 10 | 33 | **Precentral_R_5** | 6r/6v | SM | 4 | 0.56 | 0.108 | 14 | 0.386 | 0,000 | -2.08 | Provincial |
| -7 | -52 | 61 | **Precuneus_L_1** | PCV/5L | Multi/SM | 4 | 0.684 | 0.323 | 21 | 2.067 | 0.896 | 0.631 | Connector |
| -17 | -59 | 64 | **Precuneus_L_7** | 7am/7pm | CON | 4 | 0.759 | 0.554 | 14 | 0.386 | 0.849 | 0.489 | Connector |
| -31 | -11 | 0 | **Putamen_L_2** | NaN | NaN | 4 | 0.564 | 0.538 | 14 | 0.386 | 0.96 | 0.826 | Connector |
| 23 | 10 | 1 | **Putamen_R_2** | NaN | NaN | 4 | 0.793 | 0.285 | 11 | -0.334 | 0.912 | 0.679 | Satellite |
| 0 | -15 | 47 | **Supp_Motor_Area_L_1** | p24pr | CON | 4 | 0.571 | 0.392 | 5 | -1.775 | 0.99 | 0.918 | Satellite |
| 1 | -62 | -18 | **Vermis_6** | NaN | NaN | 4 | 0.508 | 0.369 | 10 | -0.574 | 0.957 | 0.816 | Satellite |

### Anatomo-functional correlates

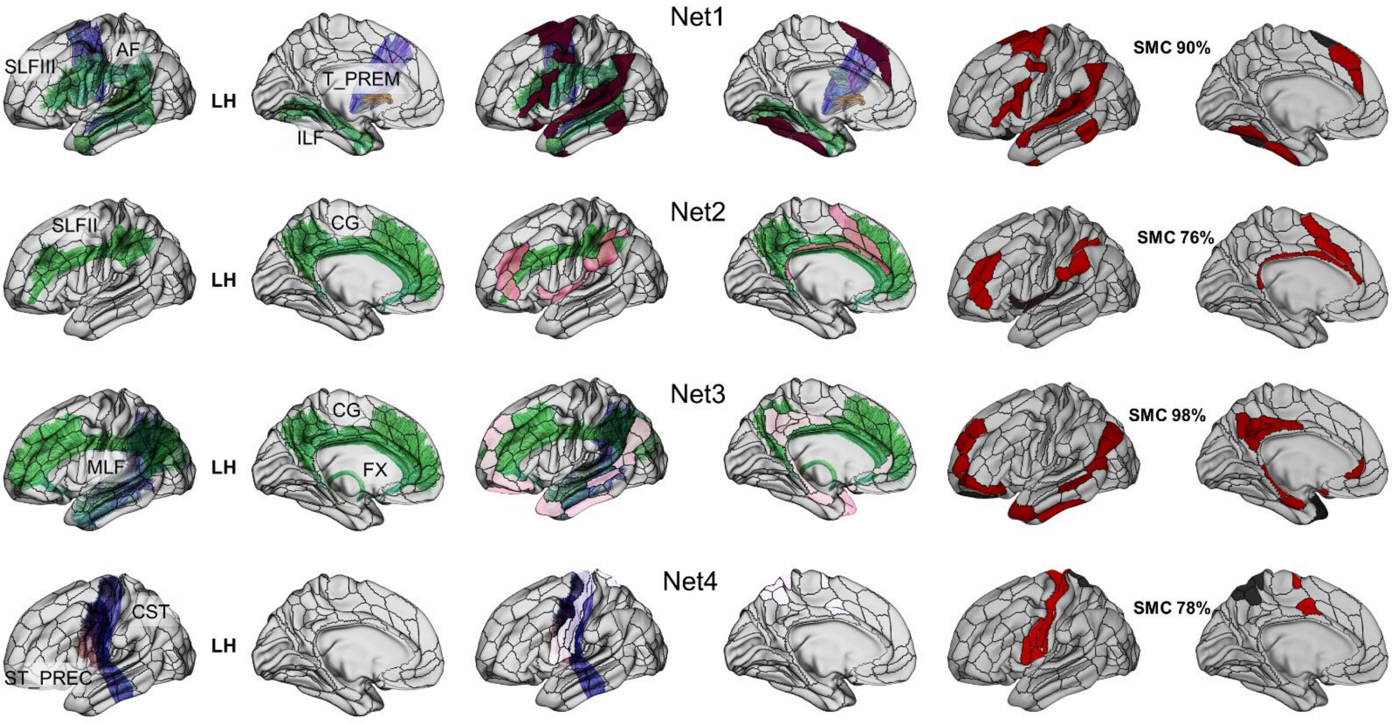

**Figure S2:** Anatomical matching (structural connectivity) with the LANG Nets.

Long-range white matter fascicles allowing a maximum spatial concordance. LH = left hemisphere; AF = left arcuate fascicle (AF); SLFIII = superior longitudinal fascicle (branch III); ILF = inferior longitudinal fascicle; T_PREM = thalamo-premotor; SLF-II = left superior longitudinal fascicle (branch II); CG = cingulum; MLF = middle longitudinal fascicle; FX = fornix; CST = cortico-spinal tract; ST_PREC = striato-precentral tract. The tracts and masks used to calculate the spatial matches are provided by TractSeg (Wasserthal et al., 2018).

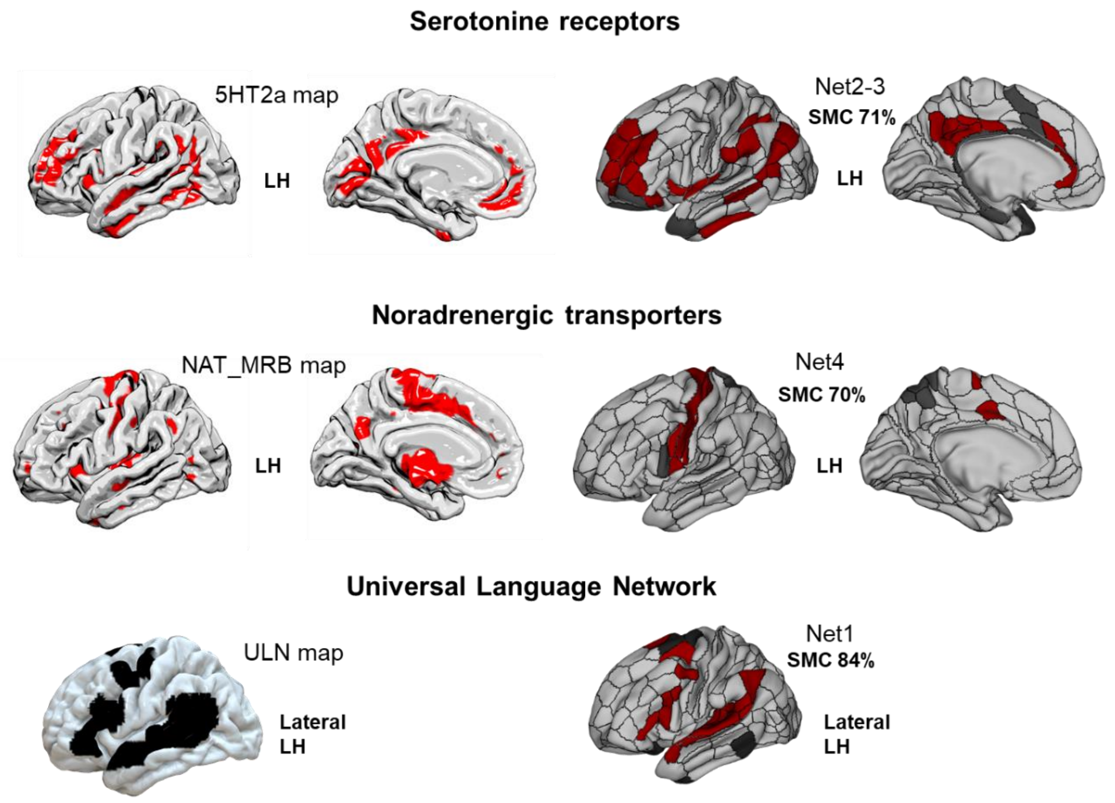

**Figure S3:** Functional matching (neurotransmitter maps) with the LANG Nets.

Significant spatial concordance with neurotransmitter receptors maps of: serotonergic (5HT2a [F18]altanserin PET; Savli et al., 2012) and catecholaminergic/noradrenergic (NET (S,S)-[(11)C]O-methylreboxetine (MRB) PET; Hesse et al., 2017) pathways. The neurotransmitters' maps are provided by JuSpace (Dukart et al., 2021). Spatial matching with the Universal Language Network functional ROIs (Ayyash et al., 2021). ULN maps are openly available here: https://osf.io/cw89s/?view_only=49981c407d784d2e88ebf6087e12fb3a

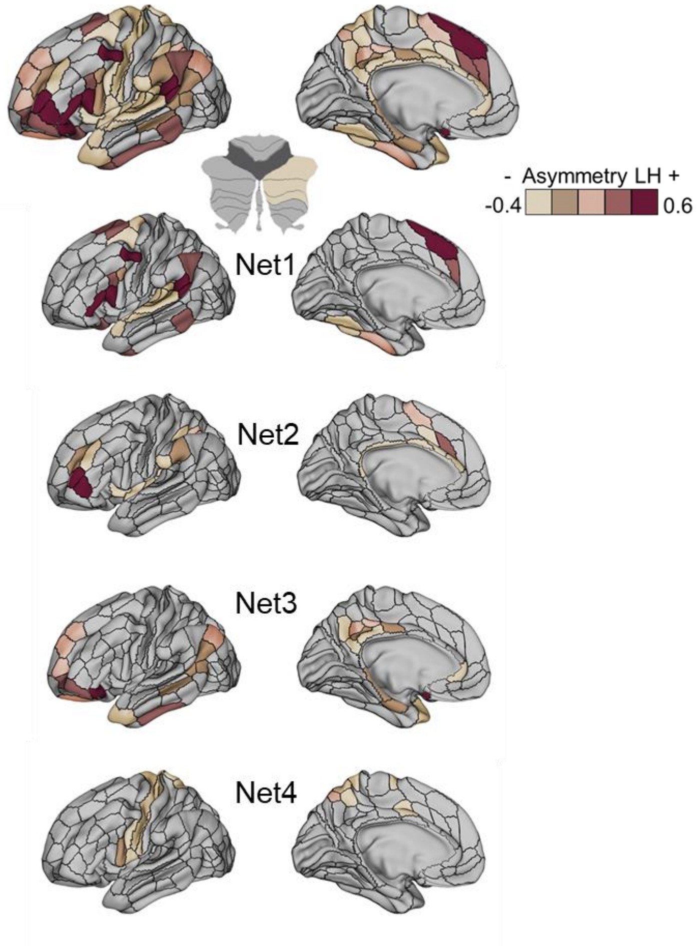

**Figure S4:** Full view (brain and cerebellum) of the functional hemispheric asymmetry across the LANG Nets.

Lateralization indexes (LIs) were calculated on all LANG regions and their homotopes, from the degree centralities (DCs). The LIs are projected here on the left hemisphere.

#### Flexibility and variability

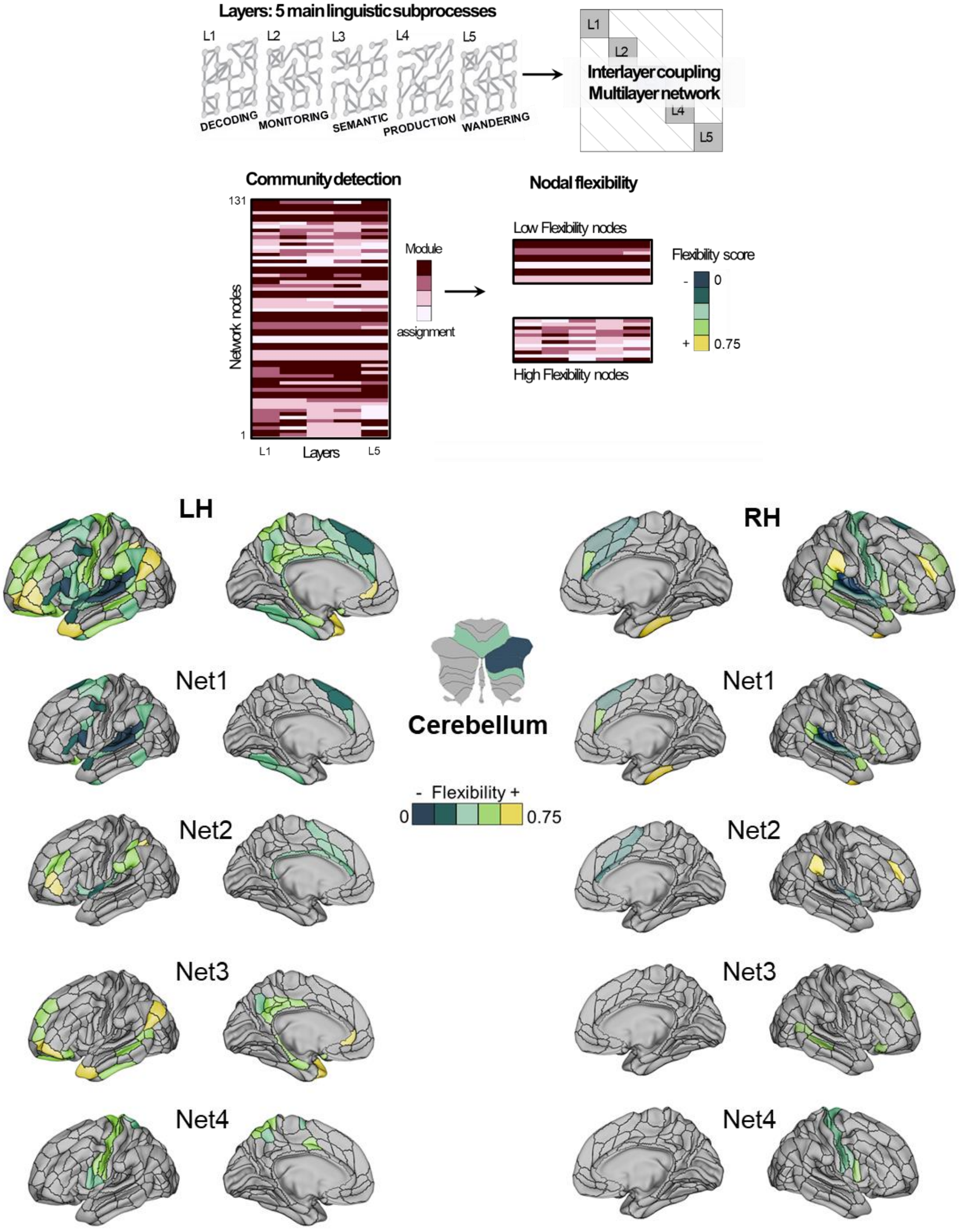

**Figure S5:** Full view (LH, RH and cerebellum) of the variability induced by the linguistic demand across the LANG Nets

We performed multilayer network analyses using a method close to that of Betzel et al. (2017). The layers of the model were constituted from the matrices of the 5 groups of tasks (i.e., latent subprocesses). To keep a common reference across the layers, the matrices were all restricted to the LANG 131 ROIs. From the communities assigned across layers, we calculated a nodal flexibility score as previously proposed by (Bassett et al., 2011).

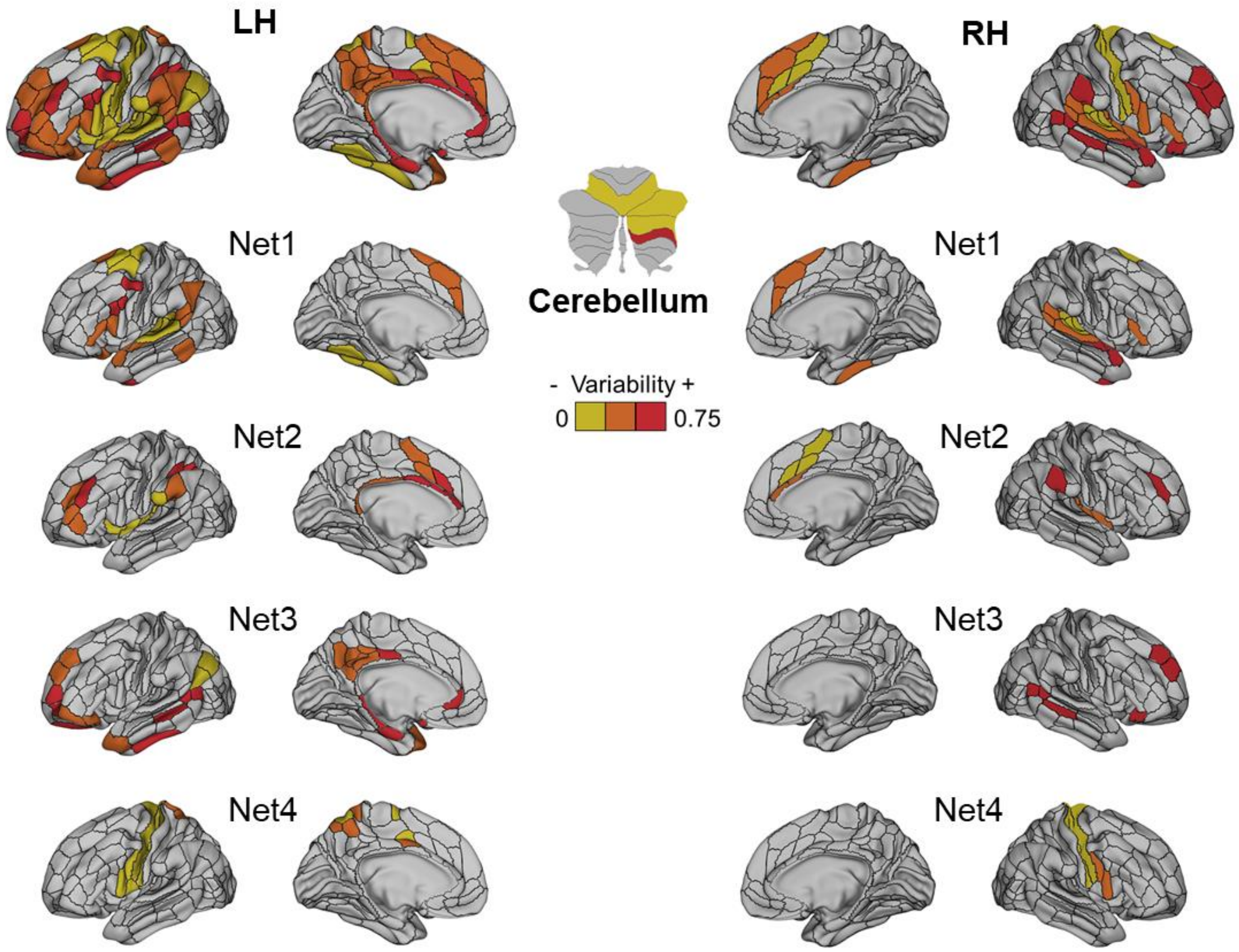

**Figure S6:** Full view (LH, RH and cerebellum) of the inter-individual variability across the LANG Nets

Inter-individual variability was estimated on the basis of fMRI signals extracted from individual maps (z scores of beta values) in the young adult cohort.

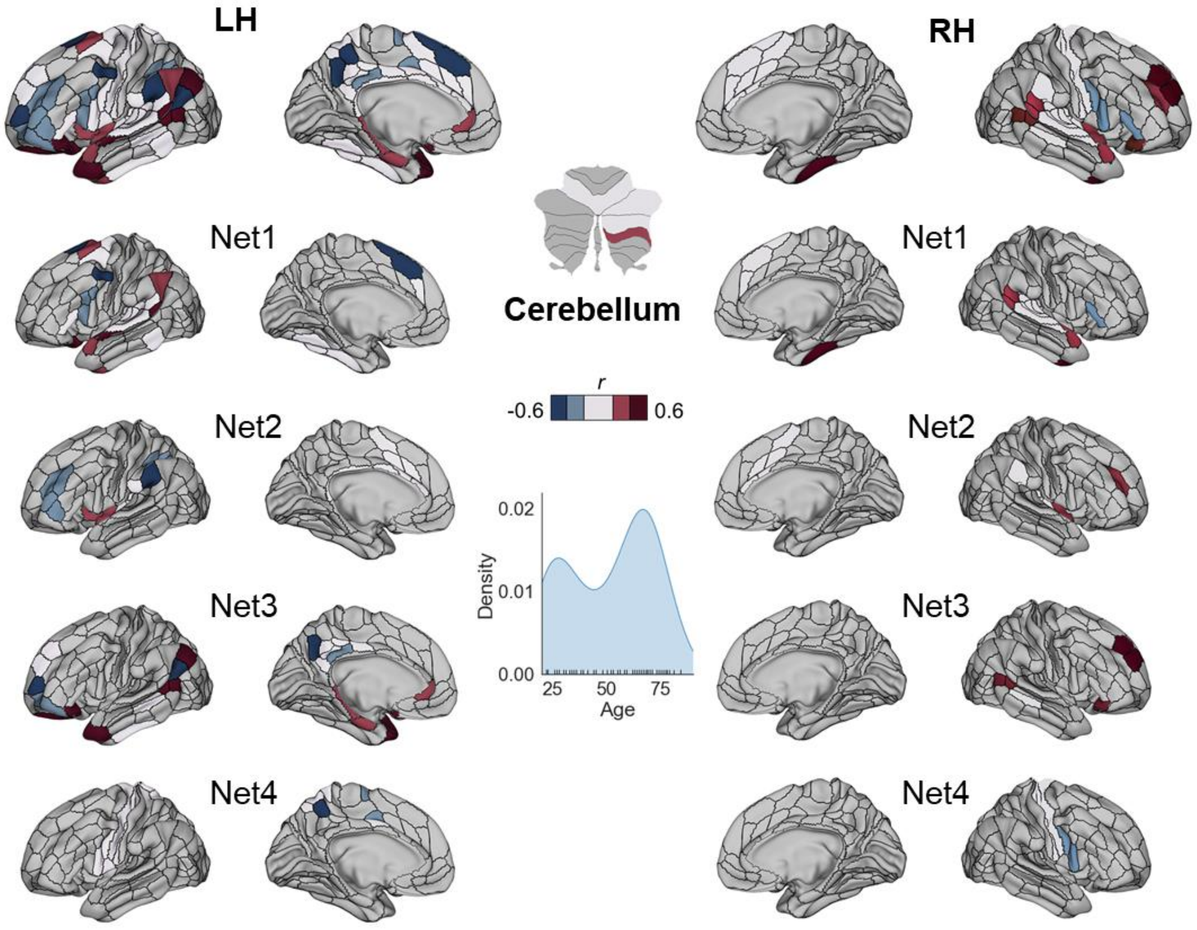

**Figure S7:** Full view (LH, RH and cerebellum) of the variability associated with age across the LANG Nets

Correlation coefficients between age and ROI DCs for each LANG ROIs. The correlations were performed on the object naming task (NAM) that includes adults over a wide range of age (n = 82; from 18 to 84 years).
